# Novel mechanisms of chemosensory adaptation to the cave environment

**DOI:** 10.64898/2026.04.03.716434

**Authors:** Natalie Choi, Edward S. Ricemeyer, X Maggs, Zicong Zhang, Masato Nikaido, Wesley C. Warren, Masato Yoshizawa

## Abstract

Smell allows animals to find food, avoid danger, and communicate through the binding of odorants to chemosensory receptors on olfactory sensory neurons. The vision-priority hypothesis predicts an antagonistic relationship between olfaction and vision, in which olfactory ability increases as visual acuity decreases along evolutionary lineages, a tradeoff that often occurs through expansion and contraction of chemosensory receptor gene families. The Mexican tetra (*Astyanax mexicanus*), a fish species with both sighted surface-dwelling and blind cave-dwelling populations, presents an ideal model for exploring the mechanisms underlying this tradeoff. Here we show that although cavefish can sense odorants at lower concentrations than surface fish, they do not have an expanded repertoire of chemosensory receptors, increased sensory neuron number or density, or enhanced expression of receptors compared to surface fish. Instead, cavefish have physiological adaptations to the olfactory epithelium, including more motile cilia and decreased flow rate through the olfactory pits. Pharmacological attenuation of flow rate in the olfactory pits in surface fish increased visits to the odorant source, suggesting that the reduced flow rate in cavefish is an adaptation leading to better foraging. This unexpected evolutionary path to enhanced olfaction as a compensation for loss of vision underscores the need for mechanistic understanding of comparative genomics.

## Introduction

The ability to detect chemicals in the environment is essential for all organisms. In animals, one of the primary forms of sensation is olfaction. Olfaction allows animals to detect the presence, concentration, and identity of chemicals in the air or water around them. They then use this information for maternal bonding, mating, kinship recognition, territorial defense, predator avoidance, and foraging^1^.

Olfactory sensitivity is highly variable and often dynamically changes during evolution. For example, among birds, archeological evidence suggests that relative olfactory bulb size, and therefore presumably olfactory ability, increased during early bird evolution but decreased during the evolution of modern birds (Neoaves)^2^. One hypothesis to explain this fluctuation is that there is a tradeoff between vision and olfaction: when vision improves, olfaction becomes less important and is therefore less consistently maintained by purifying selection; conversely, when vision deteriorates, olfaction becomes more important and is therefore targeted by positive selection (*e.g*., trichromatic color vision vs. pseudogenized olfactory receptors in primates)^3,4^.

This vision-priority hypothesis is supported by experiments in *Drosophila melanogaster* in which rearing populations in full darkness for 1,500 generations caused their optic lobes to shrink while their antenna lobes grew; these changes were reversed after only 65 generations of returning to light^5^. A similar inverse correlation between vision and olfaction across the genus *Drosophila* also supports an antagonistic relationship^6^. Conversely, primates present a counterexample to this hypothesis: the development of trichromatic vision (three color opsins) in Old World monkeys was not accompanied by a loss of chemoreceptors^7^. Thus, vision-olfaction sensory evolution may be a mosaic rather than strictly antagonistic (*e.g*., ^8^), possibly depending on their evolutionary context. If so, these sensory trade-offs would be more prominent in drastic cases, such as loss of vision over thousands of years.

To investigate the vision-olfaction trade-off and its advantages, the blind Mexican tetra presents an ideal natural experiment for uncovering the mechanisms through which a vertebrate can increase olfaction to compensate for a loss of vision. This species of teleost fish, *Astyanax mexicanus,* consists of surface-dwelling sighted populations (surface fish) and cave-dwelling blind populations (cavefish). Cavefish evolved from surface-dwelling ancestors 20-190 thousand years ago^9–11^. The cavefish adapted to dark and food-sparse environments with morphological^11–23^, metabolic^24–29^, and behavioral changes^30–40^. Notably, their vision has regressed^41–44^ while their olfactory sensitivity has increased by orders of magnitude as measured by their responses to different concentrations of amino acids^17,45,46^. In cave populations, pseudogenization of the *opsin* gene, which encodes a crucial light-sensing receptor (also a 7-transmembrane G protein-coupled receptor), has been positively selected for^47^. Together, these results may support the vision-priority hypothesis in *A. mexicanus*. However, the adaptations giving rise to the enhanced olfaction of blind cavefish remain unknown.

One classic route to investigate the evolution of olfaction is to characterize the expansion and contraction of chemosensory receptor gene families, such as the olfactory receptor (OR) gene family^4^. ORs are 7-transmembrane G protein-coupled receptors expressed in olfactory sensory neurons (OSNs) in the olfactory epithelium (OE), where binding of odorants to receptors triggers an action potential^48^. The OR gene family is among the largest in vertebrates^49^. Increased numbers of functional OR genes are thought to allow enhanced olfaction through combinatorial receptor binding^50^; for example, mice, which rely more heavily on olfaction than primates, have more total OR genes (∼1,000) than primates (∼400) and fewer OR pseudogenes^51^. Olfactory receptor (OR) repertoires in cats are smaller than those in dogs, reflecting another example of chemosensory tradeoffs, as cats rely less heavily on olfaction to hunt compared with dogs^52^.

We here show that, contrary to our expectations, independent blind populations from three different caves do not have increased numbers of functional chemoreceptor genes compared to surface fish, nor do they have increased numbers of olfactory sensory neurons or expression of chemoreceptor genes. Instead, cavefish have increased thickness of motile cilia on their olfactory lamellae, which have randomized rather than unidirectional stroking orientation, possibly contributing to decreased flow rates and increased dwell time through the olfactory pit. This unpredicted evolutionary path to compensate for the loss of vision demonstrates an essential need for a better mechanistic understanding of sensory evolution across the tree of life. Also, our study indicates possible alternative therapies via motile cilia-based treatments for patients with loss of olfactory sensation, such as those with Alzheimer’s and patients recovering from viral infection.

## Results

### Blind adult cavefish have enhanced chemosensory responses compared to surface fish

To gauge differences in olfaction between adult cave and surface fish, we tested chemosensory behavior in adults older than 2 years (*cf*. prior studies performed on juveniles 1-1.5 months old or less^17,45,46^ or groups of adults^53^, which can be subject to biases due to group behavior).

After acclimating the fish in the test tank for 7 days, 3 mL of solvent (phosphate-buffered saline: PBS) or alanine in PBS was gently injected through a tube nozzle submerged in the water, avoiding dropping sounds (Figure 1A). Alanine is highly contained in the α-helix and may signal injured prey or other food sources. Fish behaviors were tracked and scored at different alanine concentrations for the durations and frequencies of visits to four zones of the tank (top corner, bottom corner, nozzle zone, and bottom zone; Figure 1A-F). In cavefish, the top explanatory variable differentiating behavior of control versus odorant-exposed fish was the frequency of visits to the nozzle zone, whereas in surface fish the top explanatory variable was the duration of time spent in the bottom zone (Figure S1A); this different response to odorant exposure between cave and surface fish is perhaps due to spatial resolution of chemosensation to locate the food^54^. By focusing on these two explanatory behaviors respectively, we found that cavefish responded to alanine concentrations as low as 10^-10^ M with increased visits to the nozzle zone (Figure 1B) but no change in swimming distance (Figure S1B), whereas surface fish only responded to 10^-8^ M or higher concentrations of the alanine injection by spending more time in the bottom zone (Figure 1F), also increasing swimming distance at 10^-4^ M alanine (Figure S1B). Surface fish did not make more visits to the alanine source (the nozzle zone) even at higher concentrations (Figure 1C). In summary, the adult cavefish responded to 100× lower concentration of alanine (10^-10^ vs. 10^-8^ M), and their locating ability was much higher (10^-10^ M vs NA) than that of surface fish.

**Figure 1.**
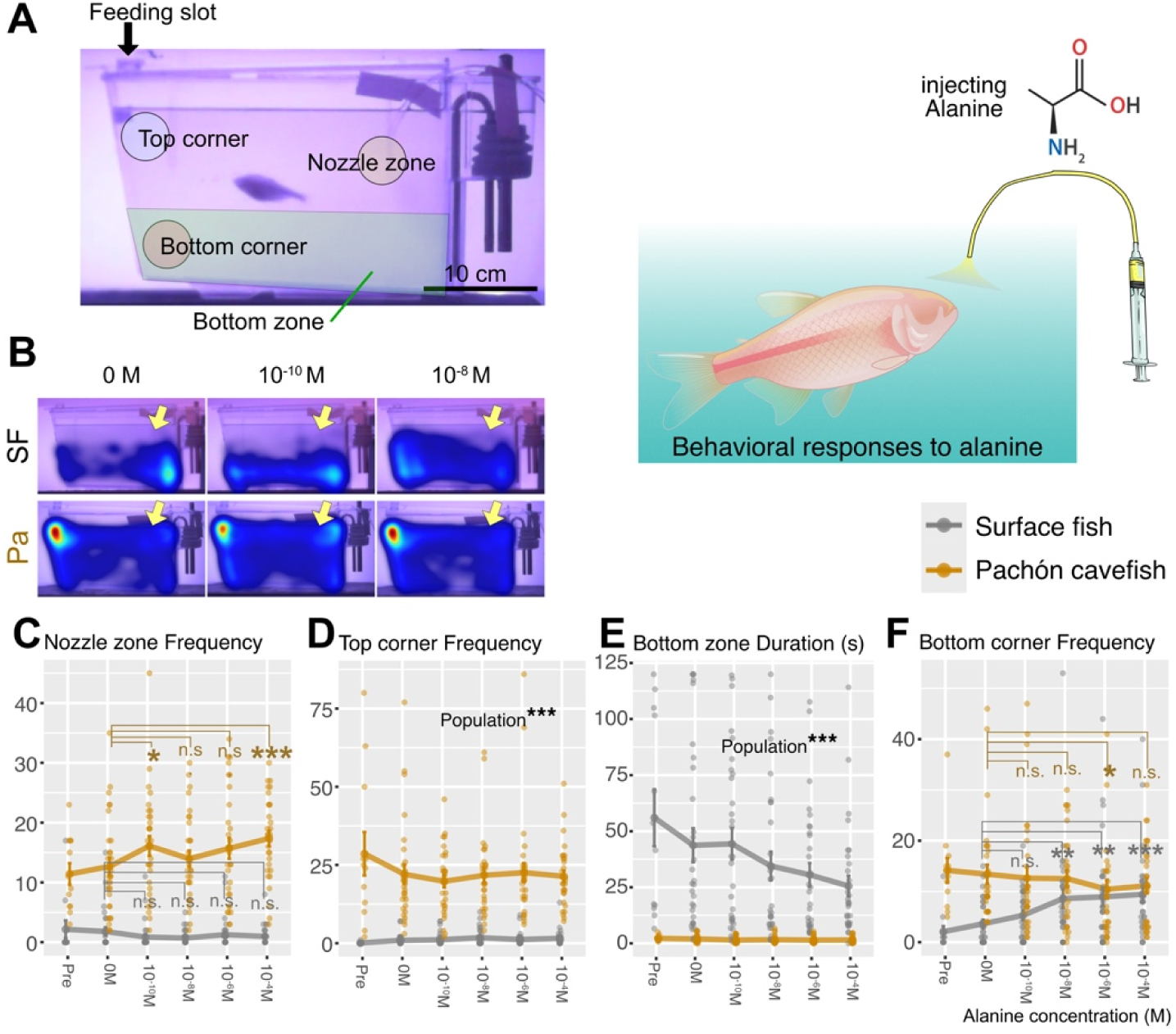
Adult cavefish were driven to the alanine odorant source while surface fish were investigated at the bottom of the fish tank. (A) Each fish was exposed to no alanine (‘pre’ treatment experiment, and injection of 3 mL ‘0 M’ alanine [no alanine] solution to the fish tank), followed by increase in concentration of alanine solution every 10 min (10^-10^ M, 10^-8^ M, 10^-6^ M, and 10^-4^ M). The left panel indicates the nozzle zone, top corner, bottom corner and bottom zone used in this study. Behavior parameters were measurements of visiting frequencies and total staying duration in each of these four zones. (B) Heat map of fish location during 10 min intervals. Nozzle zones where the alanine solution was injected are indicated with yellow arrows. Cavefish visits to the nozzle zone were more frequent (cyan) in the 10^-10^ M alanine treatment than in 0 M. (C) Quantified visit frequency to the nozzle zone (∼4 cm in diameter) during the first 6 min of the 10 min recording. Cavefish increase the visits in response to the 10^-10^ M alanine injection compared to 0 M (X^2^(1) = 7.8, P = 0.01605), while there is no detectable change in surface fish (see Data S6 for detailed statistical scores too). (D) Visit frequency at the top corner (∼ 4 cm diameter area; A) in the first 6 min. No detectable change compared with 0 M alanine in either surface or cavefish. Cavefish visited significantly higher frequency than surface fish (X^2^(1) = 64.9, P = 7.798 × 10^-16^; see Data S6). (E) Staying duration at the bottom zone in the first 6 min. No detectable change compared with 0 M alanine in either surface or cavefish. Surface fish stayed significantly longer than cavefish (X^2^(1) = 13.5, P = 2.326 × 10^-4^; see Data S6). (F) Visit frequency at the bottom corner (∼ 4 cm diameter area; see panel A) in the first 6 min. Surface fish visited significantly more in the higher alanine concentration while cavefish visited less in the higher concentration (10^-6^ M) (see Data S6). *: P < 0.05, **: P < 0.01, ***: P < 0.001. Further behavior scores and heatmap examples are in Figure S1.

### Phosphorylated ERK indicates a faster and more intense response to odorant in cavefish olfactory epithelia (OE)

To assess neuronal activity in the olfactory lamella in response to alanine treatment, we monitored an immediate early gene, phosphorylated ERK (pERK), whose signal intensities correspond to calcium-influx events involved in neuronal activation within 3 min^55^ (Figure 2A). For the latency of the pERK response, we snapshotted it at 0, 3, 5, or 10 min after bath application of 10^-6^ M of alanine, to which cavefish and surface fish are known to respond^17,53,56^ (Figure 2B-D). The pERK signal intensity per responding cell was greater in cavefish at 3 min (P < 2.2 × 10^-16^; R^2^ = 0.270 between surface and cavefish at 3 min; Figure 2B-C), and the number of the pERK positive cells was higher in cavefish (Figure 2D). For concentration response, fish were again individually treated with 0, 10^-10^, 10^-8^, or 10^-6^ M of alanine (bath treatment), and pERK-positive cells were quantified. Alanine induced pERK signals at concentrations as low as 10^-10^ M in comparison with the 0 M treatment in both surface fish and Pachón cavefish (Figure 2E, F), but the responses at 10^-10^ M in surface fish were less consistent between individuals compared with cavefish (between 0 and 10^-10^ M: P-value was 0.03037 while very low P-value, < 2.2 × 10^-16^, in cavefish; Figure 2E). In summary, these results support the behavioral differences between surface fish and cavefish: cavefish cells were activated more quickly (3 min) and consistently at 10^-10^ M than those of surface fish, guiding cavefish to the alanine source at low concentration.

**Figure 2.**
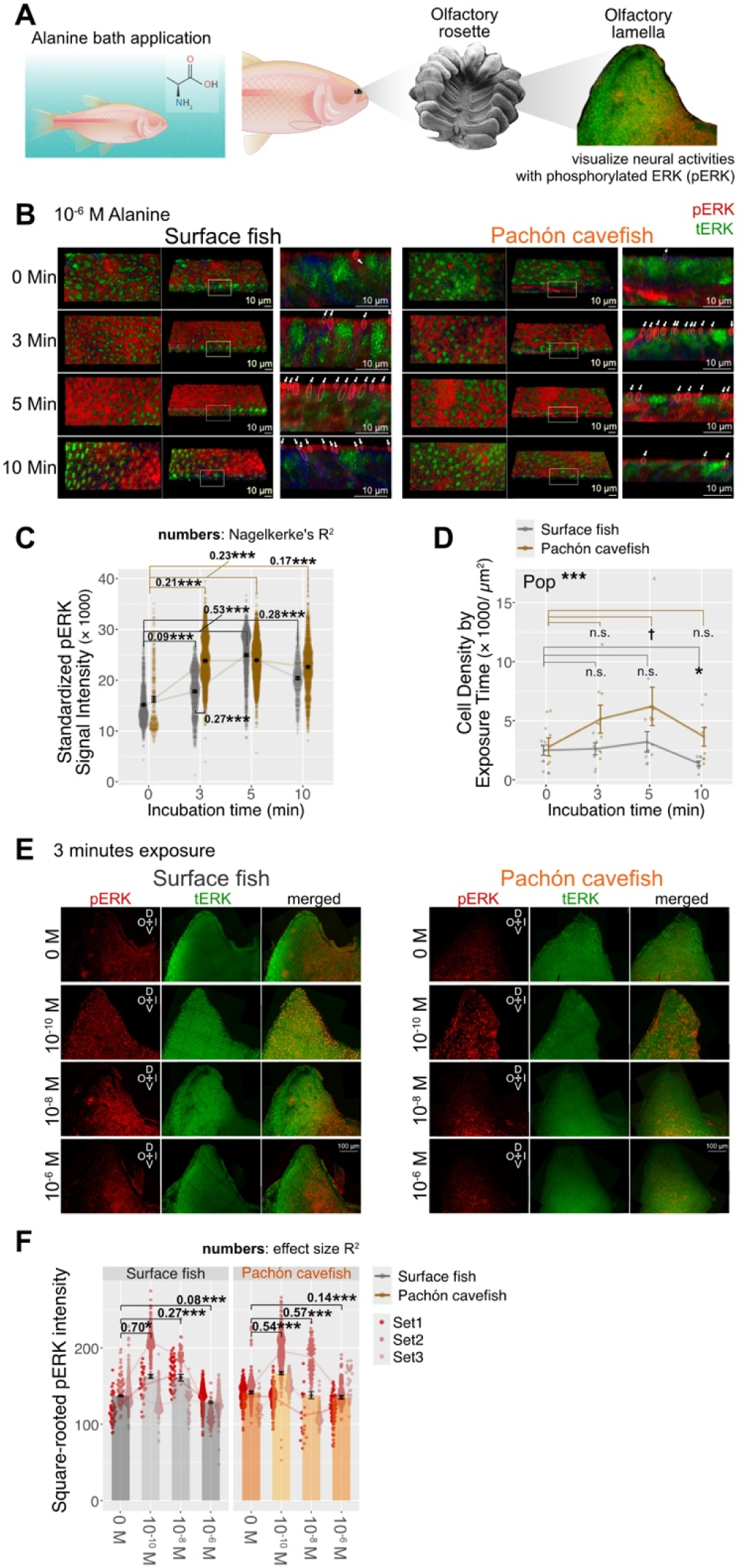
Neural activities visualized with phosphorylated ERK revealed higher alanine sensitivities in the cavefish olfactory lamella. (A) fish were bath-treated with Alanine for indicated duration (time series) or indicated concentrations (concentration series), followed by dissection of olfactory lamella on ice. (B) Examples of time snap shots of 10^-6^ M alanine exposure: 0, 3, 5, and 10 min incubation of alanine. The responded dendrites were enclosed and arrowed. Red: phosphorylated-ERK (pERK), Green: total-ERK (tERK) proteins. Total ERK (green) indicated no detectable unevenness in tERK localizations. (C) pERK signal intensity standardized with the channel gain of the confocal scanner. Cavefish cells responded to the alanine on shorter 3 min exposure than surface fish (5 min). Numbers on brackets are Nagelkerke’s R^2^ in non-linear model. (D). Cell density of pERK-positive responding cells according to the alanine exposure time. Cavefish cells responded significantly more than surface fish cells, but time differences were undetectable. The responding cells were fewer at 10 min in surface fish. (E) Examples of pERK (red) response to the alanine concentration series (0 M to 10^-6^ M, 3 min bath application per each treatment). Total ERK (green) was also labeled to observe any asymmetric expression within the lamella. There was no detectable unevenness in tERK localizations. (F) Summed pERK intensity per each detected cell was standardized with the channel gain of the confocal scanner. Square-rooted intensity scores were plotted, and compared with the pERK intensity at the 0 M alanine exposure. N = 3 individuals per treatment were used. Cavefish robustly responded to 10^-10^ M alanine. Numbers on brackets are Nagelkerke’s R^2^ in non-linear model. †: P < 0.05 before the post-hoc correction but P > 0.05 after it, *: P < 0.05, **: P < 0.01, ***: P < 0.001.

### Loss of vision is not accompanied by chemosensory gene family expansion

By addressing the enhanced olfactory ability of adult cavefish compared to surface fish, we next sought to understand its mechanistic underpinning by tracking chemoreceptor evolution, quantifying expression of these receptors in olfactory sensory neurons (OSN), and characterizing the olfactory rosette morphology and fluid flow (Figure 3A).

**Figure 3.**
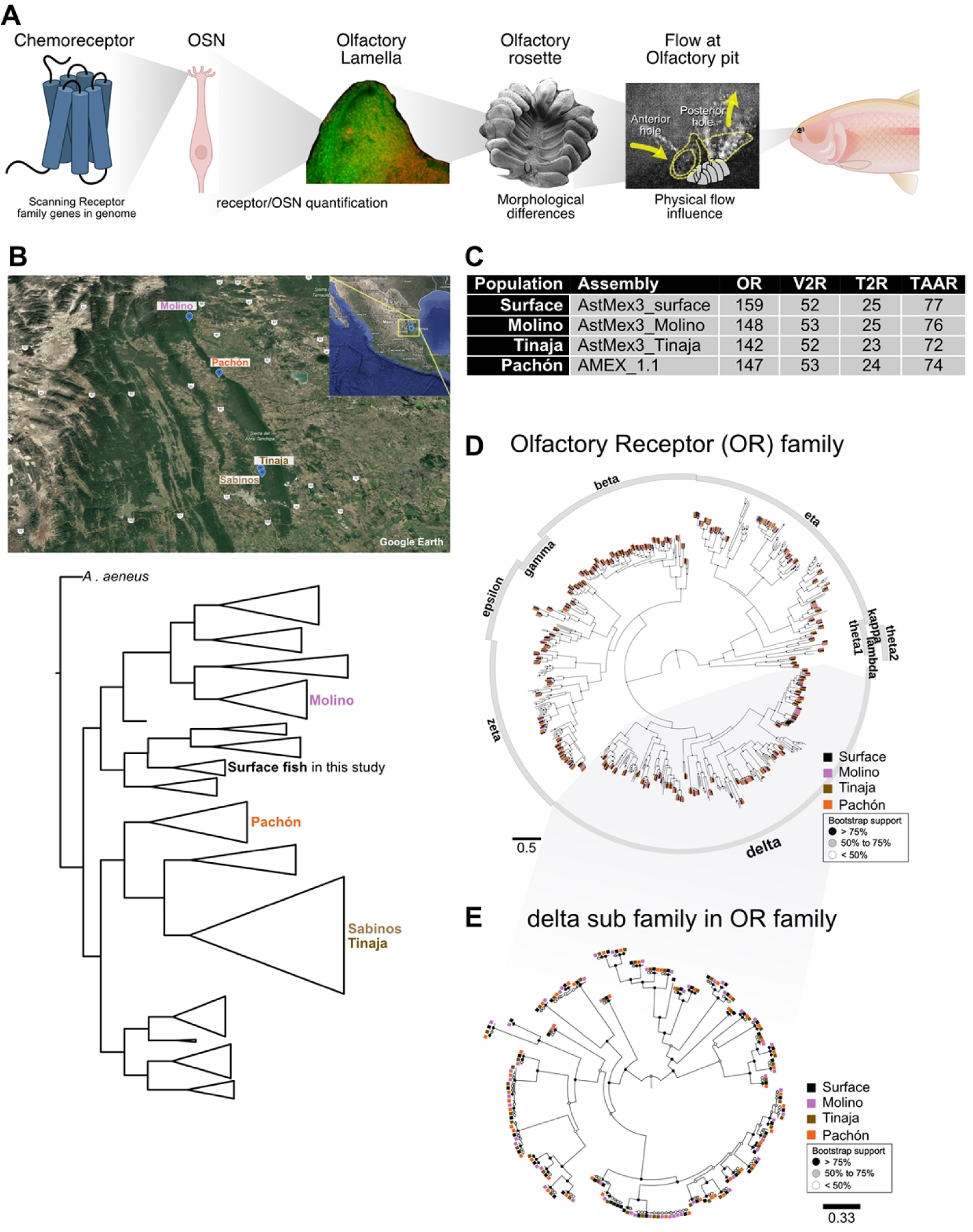
Phylogenetic tree of olfactory receptors indicated no major change in olfactry receptor family in cave populations. (A) Scheme of following studies. Analyses of chemoreceptor evolution, followed by quantifications of olfactory sensory neurons and chemoreceptors, morphological analyses and flow rate measurements. (B) Locality of the caves where the cave populations in this study were from (San Luis Potosí and Tamaulipas, Mexico), and phylogenetic tree of these populations (modified from ^11^). (C) The number of four functional chemoreceptor gene families. OR: olfactory receptor, V2R: Vomeronasal type 2 receptor, T2R: Taste receptor type 2, TAAR: Trace amine-associated receptor. (D) Phylogenetic relationship between functional olfactory receptors (ORs). (E) Phylogenetic tree of the delta subfamily of ORs. The delta subfamily is the most variable in chemoreceptor families in *A. mexicanus,* yet these expanded or reduced family members between populations were not significant compared with variation within populations. Phylogenetic trees for TAARs, V1Rs, and V2Rs are available in Figures S2, S3.

Chemoreceptor gene families contain numerous closely related (or identical) sequences, often clustered together in the genome, making them difficult to assemble accurately. Modern higher-quality genome assemblies^57^ allow better characterization of chemoreceptor gene families^49,58^. To test whether antagonistic evolution between chemoreceptors and opsin receptors occurred during cave adaptation, we annotated the intact chemoreceptors in the latest highly contiguous genome assemblies of the surface and independent cave populations (Molino, Tinaja, and Pachón cavefish) of *A. mexicanus* (Figure 3B)^59,60^. We classified genes as OR (olfactory receptors), V2R (Vomeronasal type 2 receptor), T2R (Taste receptor type 2), or TAAR (Trace amine-associated receptor)^61–64^ (Figure 3C). V1Rs and T1Rs are less diversified in teleost lineages, including *A. mexicanus,* and therefore are not listed. Our collated comparison of chemoreceptor families showed no detectable expansion in any of the three independent cave populations compared with the surface fish population (Figure 3C-E, Figures S2 and S3).

Together, these findings suggest that vision loss and opsin pseudogenization^47^ were not accompanied by chemoreceptor gene family expansion during the independent evolution of three blind cave populations, challenging the hypothesis of antagonistic evolution between visual and chemosensory systems.

### No increase in the number of olfactory sensory neurons (OSNs) in cavefish

Another alternative explanation for enhanced chemosensation in cavefish is that cavefish have adapted to darkness via a larger number of olfactory sensory neurons on the olfactory lamellae. To investigate, we used the well-established olfactory neuron markers Gα_olf_ and Gα_o_ via immunohistochemistry, both G-proteins that form the signaling complex with chemosensory G protein-coupled receptors such as OR, TAAR, V2R, and T2R^65^. Gα_olf_ is a marker for ciliated OSNs, and Gα_o_ is a marker for microvillus OSNs. We also used an antibody for Transient Receptor Potential Cation Channel subfamily C member 2 (TRPC2), another marker for microvillus OSNs, and S100, a marker for crypt neurons, to detect these neuron types^65–67^.

In the cross-sections, Gα_olf_ signals were located at the most apical cilial region (tip of dendrite) whose cell bodies located around the basal, whereas TPRC2 signals appeared in the scattered apical cells (Figure 4A). Most Gα_olf_ signals were not co-stained with phalloidin (actin) (Figure 4A), consistent with ciliated OSNs^65–67^. TRPC2-positive cell bodies were in the intermediate layer (Figure 4A), consistent with where microvillus cells locate^65–67^. Gα_o_-positive cells (microvillus OSN) were costained with phalloidin (F-actin) (Figure 4B). Their cell bodies were in the intermediate layer, consistent with TRPC2 signals (Figure 4B; *c.f.* Figure 4A), characteristic of microvillus OSN. S100-positive signals were located at the most apical, and co-stained with phalloidin, indicating crypt neurons (Figure 4B)^65,66,68^, although some S100 signals were observed in microvillus cells (Figure 4B). There were no detectable differences in the numbers or distributions of Gα_olf_-positive cells between surface fish and cavefish (Figure 4C).

**Figure 4.**
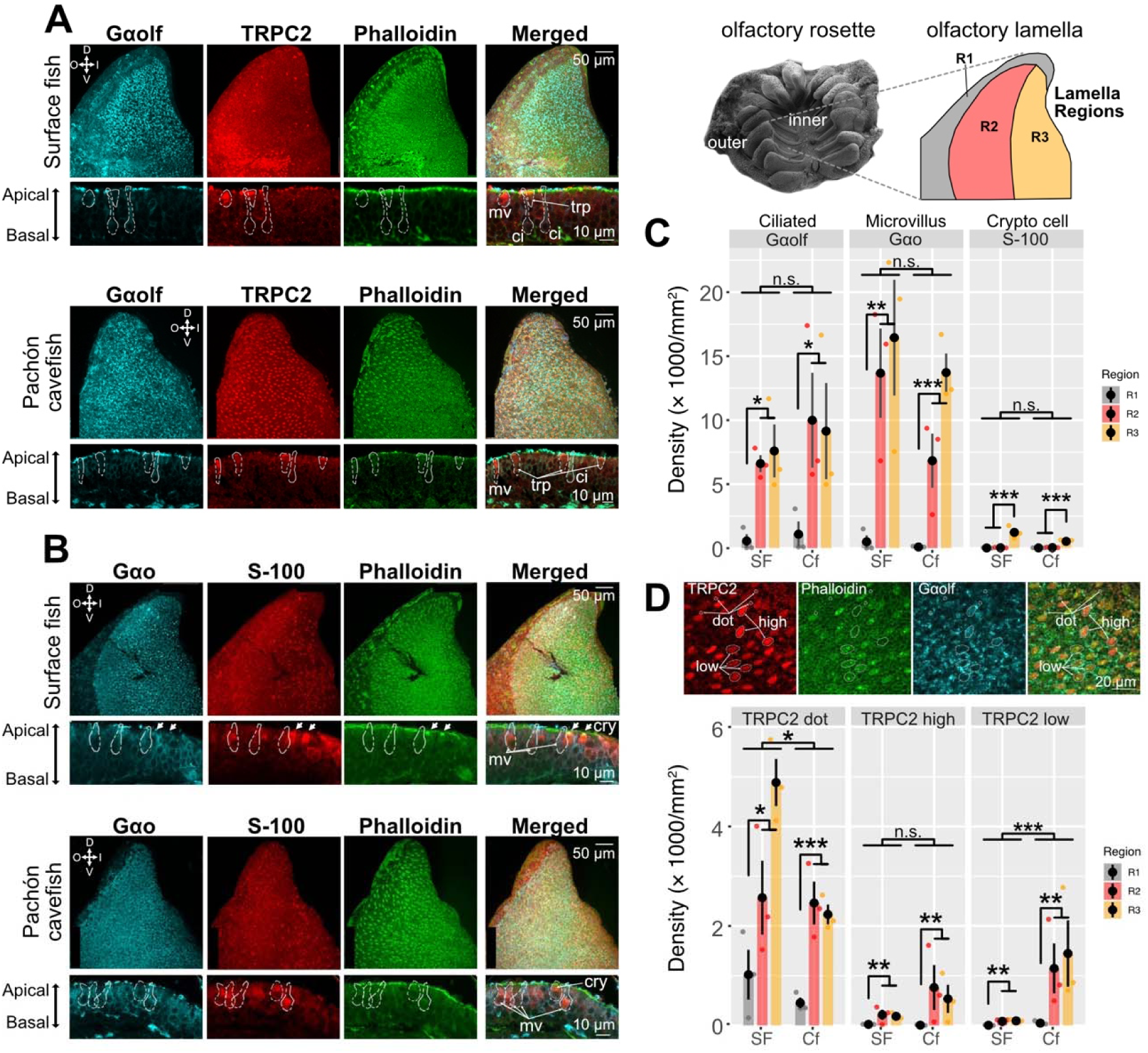
No significant increase in ciliated or microvillus olfactory sensory neurons, or crypt cells in the cavefish olfactory lamella. (A) Costained confocal images of Gα_olf_ (marker for ciliated-olfactory sensory neurons: OSN), TRPC2 (micrivillus OSN), and phalloidin (F-actin). The top row indicates the representative image of a whole lamella, where the dorsal (D), ventral (V), inner (I), and outer (O) sides within the olfactory rosette are shown. The apical-basal transverse sections of the lamella were shown below each whole lamella image. The top two rows are surface fish, and the bottom two rows are for Pachón cavefish. ci: ciliated OSN, mv: microvillus OSN, trp: TRPC2-positive cells. (B) Costained confocal images of Gα_o_ (marker for micovillus OSN), S-100 (crypt cells), and phalloidin (F-actin). Arrows indicate phalloidin and S-100-positive crypt cells. mv: microvillus OSN, cry: crypt cells. (C) Cell density of Gα_olf_, Gα_o_, and S-100 positive cells in each lamella region (R1, R2 and R3). No detectable difference between surface fish and cavefish. The R1 region contained significantly less OSNs and crypt cells. S-100-positive crypt cells were populated at R3 region at the most. (D) TRPC2-positive signals were found in the dendrites of microvillus OSN (dot), and in the unpredicted large areas (∼40 µm^2^ areas; high and low at the top panel; a cavefish example). The dot signal was significantly less in cavefish. These large-area stainings were observed only at the most apical side of the cells (*e.g*., panel A). Their lower-intensity signals colocalized with F-actin signals (top row; note that TPRC2-high signals did not colocalize with F-actin). The TRPC2 low-intensity cells were more abundant in cavefish. *: P < 0.05, **: P < 0.01, ***: P < 0.001.

We also found no detectable difference in the numbers or localizations of Gα_o_-positive and S100-positive cells, either (Figure 4C). We thus found no differences between the number of ciliated or microvillus OSNs or crypt cells in the olfactory lamella of surface fish compared to cavefish.

We regionalized the olfactory lamella as R1 (outer layer, many motile cilia but fewer OSNs), R2 (intermediate layer) and R3 (inner layer) (Figure 4). There are few OSNs in R1, and most crypt cells are in R3, but we found no difference in these distribution patterns between surface and cavefish (Figure 4C). Besides these OSN cells, TRPC2 signals exhibited large areas (∼ 40 µm^2^) with high and low levels of staining, as well as dot pattern staining (Figures 4A, 4D). The dot pattern of TRPC2 staining, representing the dendrites of microvillus OSNs, was greater in surface fish, whereas the large-area low-level TRPC staining was higher in cavefish (Figure 4D). No difference in high-level TRPC staining was detected (Figure 4D).

Together, these results indicate that although there are some differences in cell numbers between surface fish and cavefish lamella (*e.g*., TRPC2 staining, see Discussion), there is no detectable difference in major OSN types that could explain cavefish’s higher behavioral response to low alanine concentration.

### Single-nuclei sequencing confirms similar OSN composition in cave and surface

To obtain a more detailed picture of the cells present in the OE and their gene expression, and how these differ between cave and surface fish, we performed single-nuclei RNA sequencing (snRNA-seq) on OE from cave and surface fish samples (Figure 5). We used supervised clustering and marker genes from fish, mice and humans^66,69–77^ to assign a cell identity to each cell (Figure 5A; Figure S4; Data S5). We found all major cell types expected in this tissue, including olfactory sensory neuron types divided into subgroups based on maturity or differentiation. Accordingly, the olfactory sensory neuron (OSN) lineage was divided into immature OSNs (iOSN), mature OSNs (mOSN), immature microvillous OSNs (iOSN-mv), mature microvillous OSNs (mOSN-mv), and two types of differentiated OSN progenitor, intermediate neural precursors (INP) and globose base cells (GBC) (Figure 5A, Figure S4). In agreement with our immunostaining results, we found no evidence that any of these neuronal cell types are present in higher proportions in cavefish OE tissue than in surface fish (Figure 5B).

**Figure 5.**
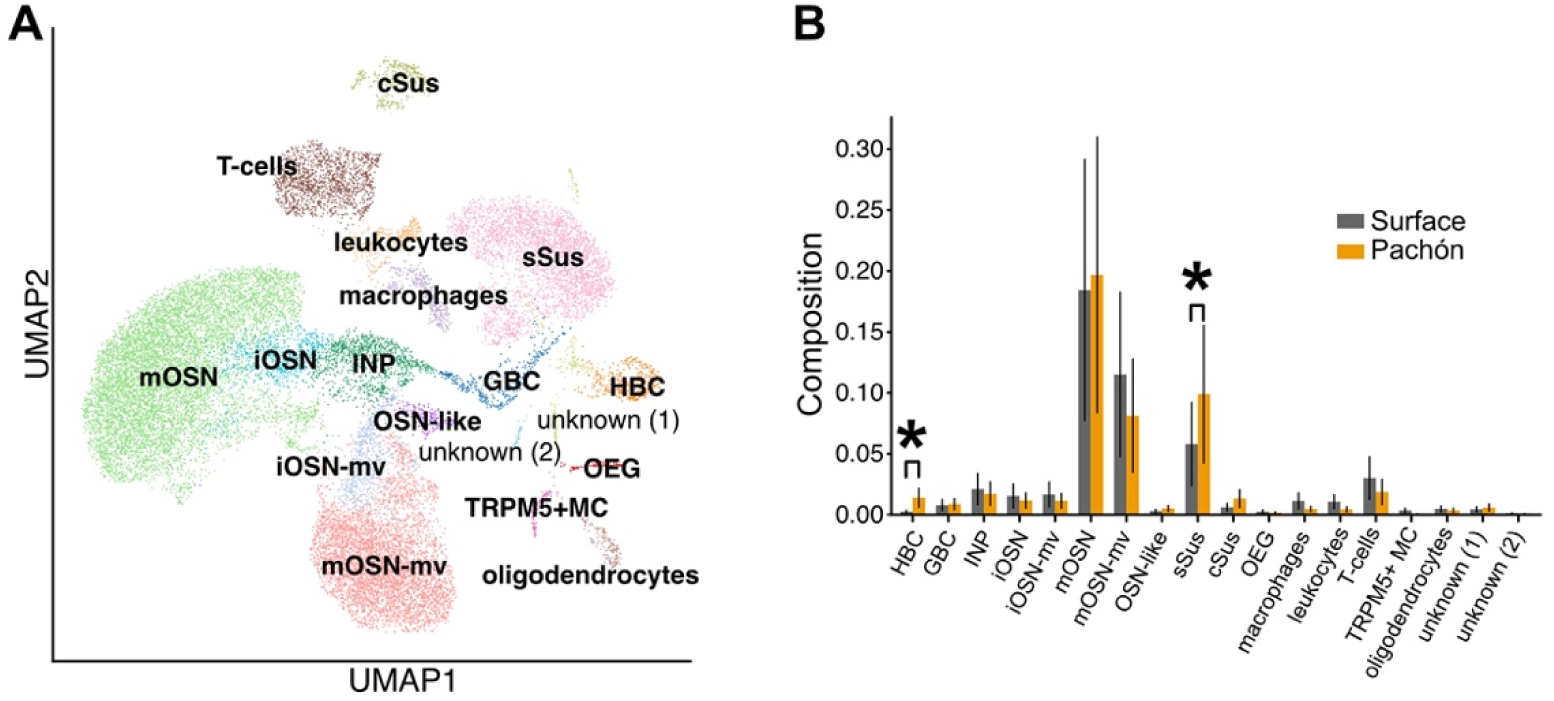
The single-nucleus RNA sequencing revealed non-significant increase in mature olfactory sensory neurons (mOSNs) in Pachón cavefish. (A) Uniform Manifold Approximation and Projection (UMAP) of the single-nucleus RNAseq results of the olfactory rosettes dissected from surface fish and Pachón cavefish. Each cell’s expression pattern was used for clustering. cSus: ciliated sustentacular cells, GBC: Globose basal cell, HBC: horizontal basal cell, INP: immediate neuronal precursor, iOSN: ciliated immature olfactory sensory neuron, iOSN-mv: micrivillus immature olfactory sensory neuron, mOSN: ciliated mature olfactory sensory neuron, mOSN-mv: microvillus mature olfactory sensory neurons, sSus: secretory sustentacular cell, OEG: olfactory ensheathing glia, TRPM5+MC: TRPM5-positive microvillar cell. See Data S5 too. (B) Proportion of cell types compared between surface fish and cavefish. Cell number propotions differed between surface and cavefish in HBC (stem cells) and sSus (secretory cells) with false discovery rate (FDR) < 0.10. *: FDR < 0.10.

However, two non-sensory cell types did show compositional differences between cave and surface samples: horizontal base cells (HBC) and secretory sustentacular cells (sSus) are both present at a higher proportion in cavefish than surface fish (Figure 5B). HBCs, identified by markers *tp63*, *sox2*, and multiple notch-family genes, are stem cells that regenerate OE tissue after it is damaged^78^ (Figure S4). sSus cells, identified by their expression of *mucin-19*, generate mucus that protects the OE and traps odorants^75^ (Figure S4). Motile-ciliated sustentacular cells (cSus, markers *foxj1a* and *dnah2*; Figure S4), on the other hand, are present in similar proportions in cave and surface OE (Figure 5B).

snRNA-seq data also allowed us to explore the distribution of receptor expression by cell type. We found that most mOSNs express only one receptor gene, whereas some cells expressed more than two receptor genes (Figure S5A)^79,80^. However, these co-expressed receptor genes are typically in the same categories expressed in the same type of olfactory sensory neurons (ciliated mOSN express OR or TAAR; microvillus mOSN expresses V2R; crypt neurons are V1R; and T2R is yet to be determined in fishes) (Figure S5B).

### No increased chemoreceptor expression in cavefish

In the absence of any compositional differences in cell counts across any class of sensory neurons, we next sought to determine whether enhanced olfaction in cavefish results from higher expression of receptor genes in similar numbers of sensory neurons. To that end, we calculated all differentially expressed genes (DEGs) between cave and surface fish samples in the mOSN and mOSN-mv clusters (Figure 6F). We found three chemosensory receptors with higher expression in the surface fish mOSN cells: two TAAR class 3 receptors (locus7241.1, locus7240.1) and one TAAR class 1 receptor (locus333.1). In mOSN-mv cells, we found one V2R sub 11 gene (locus7240.1) with higher expression in surface fish. No other chemoreceptors were differentially expressed, and no chemoreceptor gene was more highly expressed in cavefish in these clusters (Log_2_ Fold change > 2 and p-values < 0.01).

**Figure 6.**
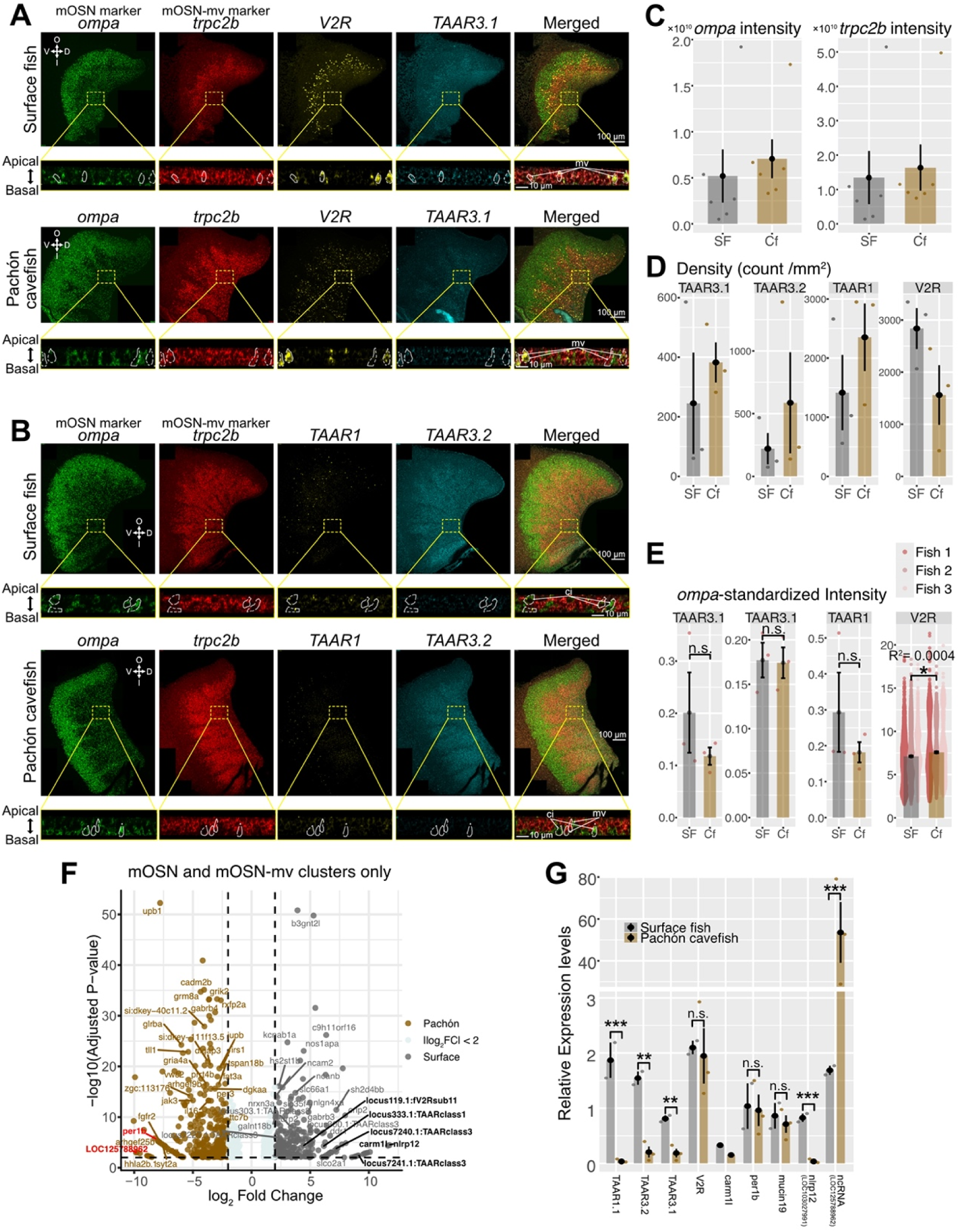
Hybridization Chain Reaction-Fluorescent *in situ* hybridization (HCR-FISH) and RT-qPCR support no significant increase in chemoreceptor gene expression in Pachón cavefish. (A) Costained confocal images of differentially expressed chemoreceptors, TAAR3.1 (locus7241.1:TAAR class3) and V2R (locus119.1:fV2R sub11), in addition to ciliated mOSN marker, *ompa*, and microvillus mOSN marker, *trpc2b*, in HCR-FISH. Dorsal (D), ventral (V), inner (I) and outer (O) sides of lamella were indicated in the inset. Transverse sections at R3 region were shown below the whole lamella images. V2R genes are known to express microvillus mOSN coexpressed with *trpc2b* but not with *ompa*, consistent with the former knowledge. *TAAR* genes are known to be expressed in ciliated mOSN while it was very difficult to thresholding TAAR3.1 expressing cells due to its weak expression level. There were no notable morphological differences in positive cells between surface and cavefish. mv: microvillus mature OSN. (B) TAAR3.2 (locus7240.1:TAAR class3) and TAAR1 (locus333.1:TAAR class1) were costained with *ompa* and *trpc2b.* TAAR3.2 and TAAR1 were weakly expressed and were difficult to evaluate positive/negative cells. Ciliated mOSN (*ompa*) and microvillus mOSN (*trpc2b*) were exclusive consistent with snRNA-seq (Figure 5, Figure S4). There were no notable morphological differences in positive cells between surface and cavefish. ci: ciliated mature OSN, mv: microvillus mature OSN. (C) Sum of signal intensity of *ompa* and *trpc2b* in an entire lamella standardised with area (µm^2^). No detectable difference between surface and cavefish, which was consistent with the snRNA-seq result (Data S1). N = 6 individuals. (D) Density (counts/mm^2^) of positive cells for each gene. No detectable difference between surface and cavefish. N = 3 individuals per group. (E) Area standardized, *ompa*-standardized intensity of chemoreceptor genes. All cells with signals were detected via the Imaris platform and the sum intensity of each of detected cells were compared (V2R). The signals of TAARs were too weak; thus, compared sum intensity per lamella (adjusted with area and *ompa* intensity). No detectable difference (TAARs) or negligible difference (V2R, Nagelkerke’s R^2^ = 0.0004) was detected. N = 3 individuals. (F) A volcano plot of differential gene expressions for ciliated mature olfactory sensory neurons (ciliated mOSN) and microvillus mature olfactory senosry neurons (microvillus mOSN) in snRNA-seq data. Data only shown for adjusted P-value < 0.01 and the absolute log2 fold change > 2.0. The genes quantified in RT-qPCR (panel G) are shown in bold characters. TAAR3.1 is locus7241.1:TAAR class3; TAAR3.2 is locus7240.1:TAAR class3; V2R is locus119.1:fV2Rsub11. (G) Top differentially expressed genes (*LOC12578862* – high in cavefish; *nlrp12* – high in surface fish), and differentially expressed chemoreceptors (TAARs, V2R), *per1b* and *carm1l* genes, in addition to a gene expressed in secretory sustentacular (sSus cells more in cavefish; *mucin 19*) were detected in quantitative reverse transcription PCR (RT-qPCR). Differential expressions in *LOC125788962*, *nlrp12*, and three TAAR genes were consistent with the snRNA-seq result (panel F). *: P < 0.05, **: P < 0.01, ***: P < 0.001.

To confirm these results and better understand the spatial distribution of gene expression in the OE, we used hybridization chain reaction fluorescence *in situ* hybridization (HCR-FISH) with probes for the four differentially expressed chemosensory receptor genes. To put the expression of these genes in the context of cell types, we also used probes for olfactory marker protein, *omp*, a marker for ciliated mOSN, and *trpc2*, a marker for mOSN-mv (Figure 6A-B). *omp* was expressed more on the ventral outer side of the olfactory lamellae, and *trpc2* was expressed on the dorsal inner side (Figure 6A-B). In agreement with snRNA-seq results, there was no difference in expression of these genes between cave and surface fish (Figure 6C).

However, unlike snRNA-seq results, HCR-FISH detected no difference in expression of the four chemosensory receptor DEGs, whether measured by density of cells expressing the genes (Figure 6D), or signal intensity standardized by *omp* expression (Figure 6E). We therefore used reverse transcriptase quantitative polymerase chain reaction (RT-qPCR) as an additional source of data (Figure 6G). RT-qPCR confirmed the higher expression in surface fish of three TAAR genes but found no difference in V2R expression.

Based on these results, we find no evidence of increased chemoreceptor expression in cavefish. Instead, we observed that expression of several chemoreceptor genes may in fact be reduced in cavefish.

### Cavefish have more cilia on OE but show slower flow through the olfactory pit

Our results in quantifying OSNs and chemoreceptors failed to explain cavefish’s ability to respond to lower concentrations of amino acids than surface fish^17,45,46^. Therefore, we next carefully surveyed morphological and structural differences in the olfactory epithelium and olfactory pits of cavefish compared with those of surface fish.

As in our immunohistochemistry, snRNA-seq, and HCR-FISH results, transmission electron microscopy (TEM) showed the expected major cell types (ciliated OSN, microvillus OSN, motile cilia-bearing sustentacular cells, and secretory sustentacular cells) in both surface and cavefish OEs (Figure 7A-D). The thickness of motile cilia was greater in cavefish than in surface fish (Figure 7A-B). Scanning electron microscopy (SEM) supports this finding, with cilia more densely covering Pachón cavefish lamellae than those of surface fish (Figure 7E). To address whether denser cilia on lamellae correspond to eyelessness, we also performed SEM on fish from three other cave populations: Molino, Sabinos, and Tinaja. We found that Molino, Sabinos and Tinaja populations have comparable cilia density to surface fish; thus, dense motile cilia specifically evolved in Pachón cavefish. In contrast, both Tinaja and Pachón cavefish have fewer lamellae per olfactory rosette (Figure S6). Despite the close relatedness of Sabinos and Tinaja cavefish (Figure 3B)^11^, their lamellae numbers are distinct (Figure 7E, S6). We also confirmed the reduced lamella number in Pachón cavefish in the non-dissected rosettes by micro-computed tomography (micro-CT) (Figure 7F).

**Figure 7.**
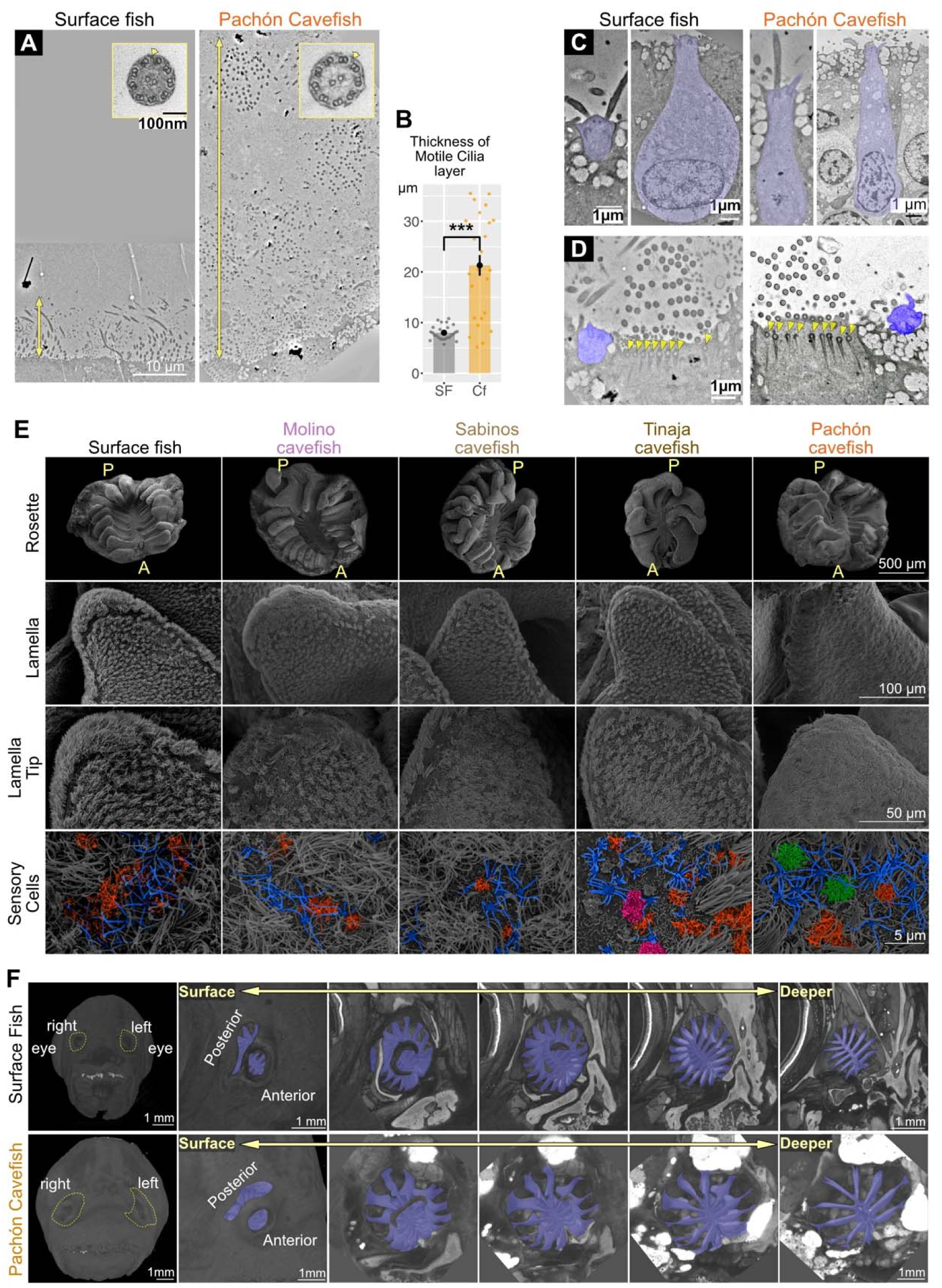
Electron microscopy and micro-CT indicated excess motile cilia with fewer lamelli evolved in Pachón cavefish. (A) Transmission electron microscopy indicated the depth of motile cilia from the base of ciliated cells. The inset indicates the cross-section of these cilia. Yellow arrows indicate the outer arm of dynein, which is, together with a 9 + 2 microtubule arrangement, a signature of motile cilia. Vertical yellow arrows indicate the depth of the motile cilia layer. (B) Quantification of motile cilia depth (see vertical arrows in panel A). (C) Examples of ciliated OSNs (blue-shaded). (D) The ciliated OSN is next to the motile cilia cells. Motile cilia cells showed basal bodies (yellow arrowheads), another signature of motile cilia. (E) Scanning electron microscopy revealed olfactory rosette and lamelli structures of surface fish and Molino, Tinaja, Sabinos and Pachón cavefish. All cavefish are blind. The lamelli of Pachón cavefish were most densely covered by motile cilia. Ciliated OSN were blue shaded, Other shade colors (red, pink and green) indicated different types of microvillus OSN. All these types of OSN are found in these five populations; therefore, no noticeable differences in OSN among these populations, consistent with the findings in Figures 4 and 6. A: anterior, P: posterior. (F) The micro-CT images support the finding of fewer lamellar numbers in Pachón cavefish than surface fish. ***: P < 0.001.

Because there are more motile cilia on the olfactory lamellae of Pachón cavefish than surface fish, we hypothesized that the enhanced olfaction of cavefish may result from more cilia promoting faster flow of water through the olfactory pits, thus exposing the OE to a greater volume of odorant in the same amount of time. To test this, we compared the flow speed in the olfactory pits between surface fish and cavefish. Surprisingly, cavefish showed slower flow than surface fish, visualized by 2 μm fluorescent microspheres (Figure 8A-B). No major attenuation in torque or speed of the motile cilia strokes between surface fish and Pachón cavefish is expected because the diameter of microtubules is indistinguishable, the outer microtubule doublets have outer dynein arms (Figure 7A, arrowheads), and the basal bodies of microtubules are indistinguishable between surface and cavefish (Figure 7D). However, the actin network supporting the microtubule basal bodies indicated significantly disturbed directions of the motile cilia orientations in cavefish (Figure S7). Thus, less-coordinated beat motions of motile cilia may underlie the slower flow rate and thus the longer retention of the odorant molecules on the lamella.

**Figure 8.**
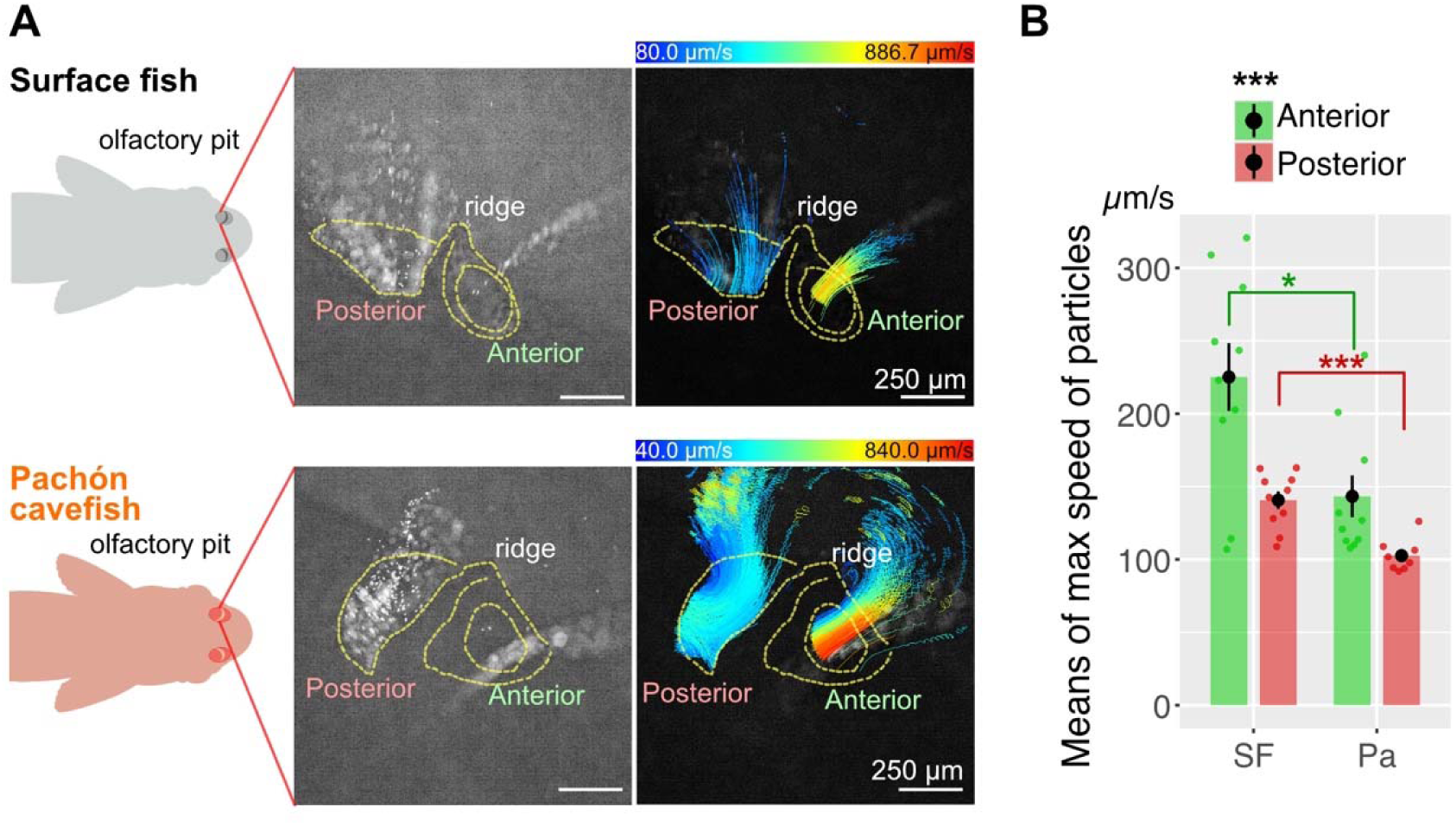
The olfactory pits of Pachón cavefish showed a slower flow rate than surface fish. (A) The left and middle panels show an imaging method of fluorescent particles (2 µm) around the nasal pits. The right panel visualized the tracked particle speeds. (B) The maximum speed of each particle was averaged in each recording session. Individual fish were recorded 3-5 sessions. Fish individuals were N = 3 per group. Statistics were calculated with the nested random effects (see Data S6). In both surface and Pachón cavefish, particle speeds at the anterior nasal pit were faster than the posterior pit, and surface fish’s flows were faster than those of cavefish. *: P < 0.05, ***: P < 0.001.

### Attenuated flow rate by a dynein inhibitor facilitates surface fish to locate the alanine source

Finally, to test whether slower nasal flow is the cause of cavefish’s ability to localize odorant source, we treated surface fish with the reversible dynein inhibitor Ciliobrevin-D at the olfactory pits (Materials and Methods, Figure S8), which slowed the nasal flow (Figure 9A).

**Figure 9.**
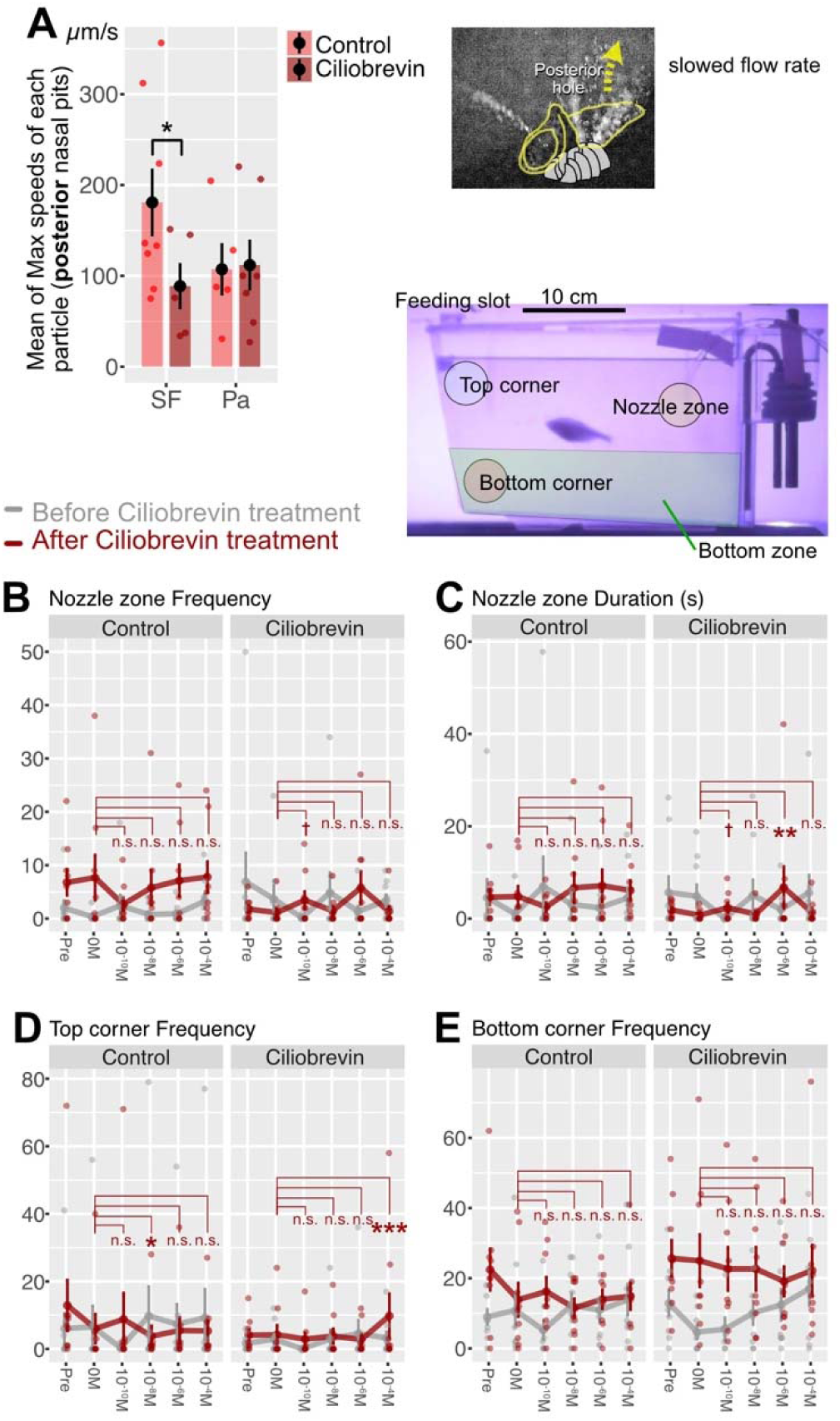
Slowed olfactory flow promoted visits to the alanine source in surface fish. (A) A 30 min treatment of a reversible dynein inhibitor, ciliobrevin-D, significantly slowed olfactory flow in surface fish. (B) Quantified visit frequency to the nozzle zone (∼4 cm in diameter) during the first 6 min of the 10 min recording. Ciliobrevin-treated surface fish tended to increase the visits in response to the 10^-10^ M alanine compared to 0 M, while no detectable change in the control or pre-ciliobrevin treatment (gray line) (see Data S6 too). (C) Visit duration to the nozzle zone in the first 6 min. Ciliobrevin-treated surface fish stayed around the alanine source of 10^-6^ M alanine (Data S6). (D) Visit frequency to the top corner (∼4 cm in diameter). Ciliobrevin-treated fish showed increased visits with 10^-4^ M alanine treatment while the control fish significantly decreased visits with 10^-8^ M alanine (Data S6). (E) Visiting frequency at the bottom corner in the first 6 min. No detectable change compared with 0 M alanine in either fish (see Data S6). †: P < 0.05 before the post-hoc correction but P > 0.05 after it, *: P < 0.05, **: P < 0.01, ***: P < 0.001.

Principal component analysis highlighted the top explanatory variables differentiating behavior of control versus treated fish as duration of time spent in the nozzle zone and frequency of visits to the bottom corner (Figure S9A), similar to those between surface and cavefish (Figure S1A). The Ciliobrevin-treated surface fish increased time spent in the nozzle zone at 10^-6^ M of alanine (Figure 9B, 9C), but not swimming distance (Figure S9B). In contrast, the control-treated surface fish did not increase frequency or duration of visits to the nozzle zone or other zones (Figure 9B-E).

From these results, we conclude that the slower flow rate in the olfactory pit enhanced the ability to locate the odorant source, likely by prolonging odorant retention on the olfactory lamellae.

## Discussion

Overall, we found an unexpected evolutionary trajectory to enhance olfactory sensation: instead of increasing olfactory sensory neurons or chemoreceptors, slowing the fluid flow in the olfactory nares that promotes the ability to locate the odorant source. We predict that our findings could suggest alternative ways to investigate the evolution of olfaction, biology-inspired chemosensing devices, and alternative therapies for loss of olfactory sensation (*i.e*., Rhinology).

In our behavioral tests, it was indicated that adult cavefish respond to the alanine in their environment better than surface fish, with cavefish increasing visit frequency at the alanine injection site compared to control, even at the lowest concentration we tested (10^-10^ M; Figure 1C). In contrast, surface fish first responded at a concentration of 10^-8^ M alanine. Because the bottom corner is where food sinks during routine feeding, surface fish individuals that detected the food smell were likely to aim for the predicted food location (*i.e*., the bottom corner and bottom zone where the food usually sits) rather than the location of the higher alanine concentration. In summary, adult cavefish responded to 100× lower alanine concentration (10^-10^ M vs. 10^-8^ M) and visited the alanine source more frequently.

It is important to note that, in our behavioral setup, the same individual fish were exposed to lower to higher alanine concentrations (see Materials and Methods) to identify the lowest alanine concentration at which the individual fish responded. Therefore, the experience of lower alanine concentrations not accompanied by actual food may have affected responses to subsequent higher alanine concentrations. Indeed, higher alanine concentrations than the lowest detectable level did not increase visits to the alanine source (Figure 1C). However, we also detected lower pERK responses to 10^-8^ and 10^-6^ M than to 10^-10^ M alanine at the cavefish olfactory epithelium (Figure 2F), where each individual fish was not exposed to multiple alanine concentrations but only one alanine concentration. Thus, decreased responsiveness at higher concentrations may be genuine rather than an artefact of the experimental design. This could possibly be explained by cavefish OSNs being tuned to detect lower concentrations of alanine so that the neuronal activities would be saturated at concentrations beyond their dynamic range, which might be induced by ephaptic inhibition—neighboring neurons inhibiting each other through an electrical field, not via synaptic connections^81,82^. This possibility should be tested in the future.

We have investigated the functional chemoreceptor families involved in olfactory sensation in teleost fishes: ORs, TAARs, V2Rs and T2R. With the latest high-quality genome, based on PacBio and Hi-C sequencing^59^, this is the first time the number of functional chemoreceptors has been effectively counted in *A. mexicanus*. The detection was performed using the FATE program pipeline, which is based on coding-region prediction, validation of predicted domains, phylogenetic placement checks, and manual validation (Materials and Methods); thus, we predict that the risk of misquantification is minimal. Small increases and decreases of receptor subtypes between surface fish and three cave populations (*e.g.,* OR delta, Figure 3E) are within the range of fluctuation per the population.

In snRNA-seq, we have annotated olfactory epithelial (OE) cells based on our best knowledge. We predict this snRNA annotation is one of the highest-quality OE cell characterizations among the non-model teleost species, supported by the high-quality reference genome^59^ (all markers and references: Data S5). Differential proportions of cell number were detected in secretory sustentacular cells (sSus) and horizontal basal cells (HBC). Cavefish’s compositional increase in sSus cells corresponded to our observation of more secretory vesicles in Pachón OE than surface fish OE in TEM (Figure 7C-D), and may be another factor in enhanced olfaction in cavefish, a possibility which should be explored in future work. HBCs are the stem cells of globose basal cells, Sus cells, and all OSN lineages^78^, so their compositional increase in cavefish could relate to a higher replacement rate of these cells in the OE to maintain high odorant sensitivity.

During TRPC2 immunohistochemistry, we observed larger area signals (∼ 40 µm^2^; TRPC2 high/low) in addition to the expected dot signals (Figure 4D). The high and low TRPC2 larger area signals were overlapped with Gα_olf_ (a marker for the ciliated OSN) and actin signals, respectively (Figure 4D), and these actin-positive low-TRPC2 cells were more numerous in cavefish than in surface fish (Figure 4D). Transverse sections of TRPC2 large signals indicated these labeled sustentacular cells (Figure 4A). Anti-TRPC2 antibody may have cross-reacted with proteins other than TRPC2, although such stainings were not reported in zebrafish and smelt^68,83^. Nevertheless, the location and stained cell shapes suggested that TRPC2-low cells are likely ciliated sustentacular cells, but further study is needed to confirm it (Figure 4A).

The fluid-flow visualization surprised us: cavefish showed slower flow around the olfactory pits than surface fish (Figure 8), despite cavefish OE having thicker motile cilia. Strikingly, pharmacological attenuation of motile cilia in surface fish enhanced their ability to locate the alanine source, suggesting that slower flow in cavefish is an adaptation that enhances olfactory sensitivity. We predict that continuous swimming, fewer lamellae, and more mucus release in cavefish further support their hypersensation of odorants.

The mystery of how cavefish have slower flow despite thicker motile cilia could be explained with two possible mechanisms. First, we observed that motile ciliary sustentacular cells on the OE seemed relatively more random in their orientation compared with those of surface fish (Figure S7). Thus, their ciliary strokes may not be coordinated to generate effective flows, allowing longer the retention time of odorant molecules in the olfactory lamellae. In addition, passive flow into olfactory pits during fish swimming replaces the water in the olfactory pits; therefore, the cavefish’s olfactory sensing does not lose time resolution, aiding in locating the source of alanine. The mechanism for capturing odor molecules may be analogous to filter-feeding animals (*e.g.,* barnacles). Through the water flow, the non-flow-aligned motions of arms/cilia capture prey/odorant and deliver it to receptors. We need to test this hypothesis in the future.

In this study, we have demonstrated that cavefish have enhanced olfaction compared to their surface-dwelling conspecifics, although this has not been accompanied by changes in the number of chemoreceptor family genes, the number of olfactory sensory neurons, or in the expression of chemoreceptors in the cavefish olfactory epithelium. Instead, evolution found a creative working solution to improve chemosensory ability: altering fluid dynamics at the olfactory epithelium through the action of motile cilia. It will be interesting to see if other independent cave populations of *A. mexicanus* as well as numerous aquatic cave-dwelling species follow this same working model. Further research promises to reveal alternative examples of how evolved olfactory sense compensates for vision loss. The knowledge gained herein suggests therapeutics for olfactory loss via changes in fluid (*e.g*., mucus layer) on the olfactory epithelium, such as to prevent the progression of Alzheimer’s (olfactory loss is the initial diagnosis^84^) and/or facilitate recovery from COVID infections (a chronic loss even after recovery^85^) could be considered.

## Materials and Methods

### Fish maintenance and rearing in the lab

*A. mexicanus* surface fish used in this study were the laboratory-raised descendants of original collections created in Balmorhea Springs State Park in Texas in 1999 by Dr. William R. Jeffery. Cavefish were laboratory-raised descendants originally collected from Cueva de El Pachón (Pachón cavefish) in Tamaulipas, Mexico in 2013 by Dr. Richard Borowsky.

Fish (surface fish and Pachón cave populations) were housed in the University of Hawai‘i at Mānoa *Astyanax* facility with temperatures set at 21 ± 0.5°C for rearing, 24 ± 0.5°C for behavior experiments, and 25 ± 0.5°C for breeding ^34,86^. Lights were maintained on a 12-h:12-h light:dark cycle ^34,86^. For rearing and behavior experiments, the light intensity was maintained at 30–100 Lux. Fish husbandry was performed as previously described ^10,34,86^. Fish were raised to adulthood and maintained in standard 42-L tanks in a custom-made water-flow tank system.

Approximately 10% of tank water are replaced 3 times per day with the conditioned fish water, which is based on the reverse-osmosis filtered water adjusted its conductivity about 700 µS with Reef Crystals Reef Salt (Instant Ocean Spectrum Brands, Blacksburg, VA, USA) and pH with Hydrochloric Acid (Thermo Fisher Scientific, Waltham, MA, USA) from the original 8.3 to 7.0.

Adult fish were fed a mixed diet to satiation twice daily starting 3 h after the lights were turned on (Zeitgeber time 3 [ZT3] and ZT9; Zeigler), including Adult zebrafish irradiated diet (Zeigler Bros, Inc, Gardners, PA), TetraColor Tropical Fish Food Granules (Tetra, Blacksburg, VA, USA), Jumbo Mysis Shrimp (Hikari Sales USA, Inc., Hayward, CA, USA), and Ken’s Ultra Intense Spirulina Flake (Ken’s Fish & Pet Supplies, Taunton, MA, USA). All fish in the behavioral experiments were between 3 and 6 cm in standard length and between 3 and 5 years old. All fish care and experimental protocols were approved under IACUC (17-2560) at the University of Hawai‘i at Mānoa.

### Behavior assay and analysis in response to alanine

To compare the differences in response to alanine injection/perfusion to the fish arena, we first individually housed fish in a 3.5-L techniplast ZebTEC fish housing tank (approximately 24 × 10 × 15 cm in length × width × height, respectively). The recording chamber was illuminated with a custom-designed IR LED source in a light-controlled room on a 12-h:12-h cycle (was turned on at 7:00 am and turned off at 7:00 pm each day). The visible light during behavior recordings was approximately 30–100 Lux. L-alanine solution (L-alanine, MilliporeSigma; 10^-10^, 10^-8^, 10^-6^, and 10^-4^ M of alanine in PBS) was supplied through a 3-way valve ([company]) attached with a 20 cm of silicone tube whose tip was submerged in the fish water in the fish housing. Adult female fish, which were raised in the 3.5 L ZebTEC tank, were acclimated to a carefully cleaned 3.5 L ZebTEC tank with the injection tube for at least 7 days before recording. Each tank was aerated. Full water changes were performed every two days and also a day before recording. Fish were fed with 5-10 particles of Zebrafish Diet (Zeigler Foods, Gardners, PA) every morning, once a day from the feeding slot (Figure 1A). The 3-way valve attached with a 20 cm tube was also attached with the syringe-1 filled with 3 mL of alanine solution or just PBS, and with the syringe-2 filled with 3 mL of the conditioned fish water. The 3 mL conditioned fish water in syringe-2 was to flush the 20 cm tube, delivering all the alanine solution to the fish tank.

Behavior was recorded after 7 days of acclimation in the following sequence: (i) 10 min recording without any disturbance, (ii) recording right after injecting 3 mL of PBS followed by 3 mL of PBS to flush the injection tube for 10 min, followed by (iii) injecting 3 mL of 10^-10^ M alanine + 3 mL PBS flush then recording fish behavior 10 min, (iv) 3mL of 10^-8^ M alanine + flush then recording for 10 min, (v) 10^-6^ M alanine + flush and the 10 min recording, and (vi) 10^-4^ M alanine + flush and the 10 min recording.

This recording session began 1–2 h after turning the light on (ZT1–2) without feeding. The air stone was removed at least 1 h before the first recording. Six fish were recorded at once by using two camera setups. The entire recording finished within 80 minutes. Videos were recorded at 15 frames/s using a USB webcam with an IR high-pass filter. Videos were captured by VirtualDub2 software with the x264vfw codec and subsequently processed using Ethovision XT (Version 16, Noldus Information Technology, Wageningen, Netherlands). The tracking parameters for detection were as follows: the detection was set to “subject brighter than background” and brightness contrast was set from 20 to 255; the current frame weight was set to 15; the video sample rate was set to 15 frames/s; and pixel smoothing was turned off. The X-Y coordinates of each fish were subsequently processed using custom-written Perl (v5.23.0, www.perl.org) and Python scripts (3.8) (https://zenodo.org/record/8137637). Water temperature was monitored throughout the recordings, and no detectable differences were observed during the light and dark periods (23.0 ± 0.5°C).

### Identification and phylogenetic reconstruction of chemoreceptor genes

Using the four *A. mexicanus de novo* genome assemblies, we identified the repertoires of chemosensory receptors (OR, TAAR, V1R, T2R, and V2R). The identification pipeline was implemented using the FATE program. Briefly, previously identified intact OR^61^, TAAR^62^, V1R^87^, T2R^63^, and V2R^64^ genes were used as queries for tBLASTn v2.14.0+ searches. The resulting hits were parsed with GeneWise v2.4.1^88^ to predict coding regions, and the predicted sequences were searched against the UniRef50 database using BLASTx (last accessed on 3 August 2022) to filter out non-homologous hits. To remove false-positive hits and classify candidates into known subfamilies, we aligned all intact candidate sequences together with previously identified genes using MAFFT v7.505^89^ with the --linsi option and inferred phylogenetic trees with RAxML-NG v1.2.0^90^ with the --all -bs-trees 100 options. The best-fitting substitution models for tree construction were selected with ModelTest-NG v0.1.7^91,92^ based on AICc scores. All sequences derived from the *A. mexicanus* assemblies that clustered with known subfamilies in the phylogenetic analyses were treated as true positives. The reconstructed phylogenetic tree was visualized by TreeViewer v2.2.0^93^

For pseudogene identification, we first extracted gene regions that had been classified as pseudogenes by GeneWise in the above procedure and that were longer than 600 bp. These sequences were then added to the corresponding multiple sequence alignments using the --add option of MAFFT. Then, we used the --tree-constraint option of RAxML-NG to place the pseudogene taxa onto the gene trees inferred from intact sequences. In the last, pseudogenes that clustered with known subfamilies under this process were regarded as true positives. Each pseudogene region was carefully examined by checking its sequence manually and by checking the their placement in the phylogenetic tree, thus we predict the chance of misquantification (fail to count or false call of an actual pseudogene as a functional gene) is very limited.

### Single nucleus RNA sequencing

#### Single Nuclei Isolation

Olfactory epithelium was flash-frozen and stored at -80 °C. Per sample, nuclei isolation was performed on a Singulator 100 (S2 Genomics) with the Standard Nuclei Isolation Protocol. The entire tissue (∼25 mg) was loaded on the isolation cartridge. After initial isolation, the solution was spun at 500g for 5 minutes at 4° C, and the supernatant was discarded. The pellet underwent two washes using 250 µL of Nuclei Storage Reagent (S2 Genomics; 100-063-405) containing 0.4 U/µL Millipore Protector RNAse inhibitor (Sigma Alrdich; RNAINH-RO). After each wash, nuclei were centrifuged at 500g for 5 min at 4°C, and the supernatant was discarded. The final pellet was resuspended in 250 µL of the wash buffer solution. Wide bore tips were used throughout. Isolated nuclei were visualized and quantified on a Cellometer K2 model (Revvity).

#### Single Nuclei Sequencing

Sequencing libraries were prepared with a Next GEM Single Cell 3 Kit v3.1 (10X genomics; PN-1000268). Library concentration and fragment size distribution were determined with the 5200 Fragment Analyzer version 3.1.0.12 (Agilent Technologies). Paired-end 100bp sequencing of sample libraries was carried out on an S4 flow cell on an Illumina Novaseq 6000 instrument at the Genomics Technology Core at the Bond Life Sciences Center (Columbia, Missouri).

#### Generating count matrices

We first prepared a custom reference based on the reference assembly AstMex3^59^, containing the annotated olfactory receptor genes. To prioritize these genes and avoid multi-mapping, we combined the gene set from the published annotation with our olfactory receptor annotations, giving precedence to the olfactory receptor annotations in cases of overlap between the two annotation sets. We then ran cellranger v7.2.0^94^ command mkref with default options to index this reference. We aligned reads from each sample to the reference and created raw cell/gene count matrices using the cellranger count command with default parameters and then filtered barcodes most likely to represent doublets or ambient signal by running cellbender v0.3.0^95^ command remove-background with default options.

#### Clustering

We next used the scanpy platform v1.9.8^96^ to perform a standard supervised clustering process on the combined set of cells from all samples. Briefly, we filtered out genes expressed by fewer than 3 cells, cells expressing fewer than 200 or more than 3,000 genes, cells with more than 5,000 total counts, and cells with 4% or more counts assigned to ribosomal genes; we then normalized and log-scaled counts, selected highly-variable genes, ran principal components analysis on the highly-variable genes, integrated the principal components by sample with harmony^97^ to mitigate batch effects, calculated the neighbor graph, and clustered the cells using the leiden algorithm with resolution 1.1.

#### Annotation

We generated a list of potential marker genes for each cluster using scanpy’s rank_genes_groups function. We manually annotated cell identities based on orthology to established cell type marker genes in other species, using orthology from Roback & X et al, (2025 submitted). Cluster assignments and the marker genes used for each assignment are in Data S5 (also, Figure S4).

After this, we merged clusters with the same cell type assignments.

#### Differential composition

We found cell types with different composition in the cave versus the surface samples using scCODA v0.1.9^98^ with false discovery rate (FDR) 0.1.

#### Differential expression

We compiled lists of genes with differential expression between cave and surface samples for each cluster using the pseudobulk method^99^. Briefly, for each sample, we summed the gene counts across all cells to produce a single vector of gene counts per sample, and then ran pydeseq2^100^ with the single design factor “population” (i.e., cave or surface). We considered a gene to be differentially expressed if the absolute value of its log2 fold change (LFC) is at least 1 and its multiple-hypothesis-adjusted p-value is less than 0.01.

Scripts and notebooks used to perform all bioinformatic analyses are available in the project code repository.

### Scanning Electron Microscopy

*A. mexicanus* olfactory epithelia were dissected and fixed with 2% paraformaldehyde and 2.5% glutaraldehyde in 0.1M sodium cacodylate buffer (pH 7.4) for 1-2 hours at room temperature. Samples were washed twice in 0.1M cacodylate buffer for a total of 20-30 minutes. Tissues were post-fixed in 1% Osmium tetroxide in 0.1M cacodylate buffer for 1 hour and subsequently dehydrated in a graded ethanol series (30%, 50%, 70%, 85%, 95%; two changes of 3 each), followed by three changes of 10 minutes each in 100% ethanol. Dehydrated samples were critical point dried in a Samdri-795 (Tousimis Research Corporation, Rockville, MD, USA), mounted on stubs with a conductive paint, and coated with Au/Pd in a LTD Hummer Sputtering System (Anatech USA, Sparks, NV, USA) for 1 minute. Prepared specimens were imaged using a S-4800 Scanning Electron Microscope (Hitachi High-Tech Corp., Tokyo, Japan). Specimens used were N = 1-2 biological replicates per morph and cave population.

### Transmission Electron Microscopy

Fixation and dehydration procedures were identical to the SEM protocol, except that samples were post-fixed with 1% osmium tetroxide following washing. Briefly, *A. mexicanus* olfactory epithelia were fixed with 2% paraformaldehyde, 2.5% glutaraldehyde, and 2mM CaCl_2_ in 0.1M sodium cacodylate buffer (pH 7.4) for 1-2 hours at room temperature. After washing, tissues were post-fixed with 1% osmium tetroxide in 0.1M cacodylate buffer for 1 hour and dehydrated through a graded ethanol series. Dehydrated samples were transitioned to propylene oxide (three changes of 10 minutes each) and infiltrated overnight in a 1:1 mixture of propylene oxide and epoxy resin. The following day, samples were transferred to 100% epoxy resin and incubated for 2–8 hours, then transferred to fresh epoxy resin for 1–4 hours, and finally to a third change of fresh resin for 0.25–1 hour. Samples were then placed in molds with epoxy resin and polymerized at 60°C for 2-3 days. Ultrathin sections (90-110 nm, gold interference color) were collected on copper grids and imaged on a HT7700 Transmission Electron Microscope (Hitachi High-Tech Corp.). Specimens used were N = 1 biological replicates per morph.

### Micro-CT

Heads from euthanized A. mexicanus (surface fish and Pachón cavefish) were dissected by removing gills. The samples were then fixed by submersion in 4% paraformaldehyde (Electron Microscopy Sciences, Hatfield, PA, USA) in 1× Phosphate-Buffered Saline (Santa Cruz Biotechnology, Inc., Dallas, TX, USA) for 18-24 hours at 4°C with vigorous shaking.

Following fixation, After washing samples with 1× PBS twice, the head tissue was stained with Lugol solution (MilliporeSigma, Burlington, MA, USA) for 48 hours at room temperature with gentle shaking for contrast enhancement. Subsequently, each fish head was embedded in 2% low-melting-point agarose (Thermo Fisher Scientific, Waltham, MA, USA) and imaged via SkyScan 1272 micro-CT scanner (Bruker Corp., Billerica, MA, USA). Raw projection images were processed into cross-sectional slices using Bruker’s NRecon software (Bruker).

Reconstructed slices were rendered using the CTvox volume rendering software (version 3.3.0, r1403, Bruker). N = 2 biological replicates per morph were scanned.

### Immunohistochemistry

The adult *A. mexicanus* were anesthetized in ice-cold water for 10 min by monitoring cessation of any gill operculum or body movements. Olfactory epithelia were dissected and fixed with fresh in 4% paraformaldehyde (PFA: Electron Microscopy Sciences, Hatfield, PA, USA) in 1× Phosphate-Buffered Saline (PBS: Santa Cruz Biotechnology, Inc., Dallas, TX, USA) for 18 hours at 4°C with gentle agitation. Tissues were rinsed with ice-cold 0.25% PBT (1× PBS and 0.25% Triton® X-100 (Acros Organics, Geel, Belgium) and incubated in a blocking buffer, which consists of 1% Dimethyl sulfoxide (MilliporeSigma, Burlington, MA, USA), 2% Bovine Serum Albumin Fraction V (Roche Diagnostics, Rotkreuz, Switzerland), and 10% of Normal Goat Serum (Vector Laboratories, Inc., Newark, CA, USA) in 0.25% PBT, for 1 hour at room temperature, followed by overnight incubation of primary antibody at 4°C with gentle agitation (see below).

The primary antibody solution consisted of mouse anti-G_αolf_ (1:500 dilution; Santa Cruz Biotechnology, Dallas, TX), mouse anti-G_αo_ (1:100 dilution; Santa Cruz Biotechnology), rabbit anti-zebrafish TRPC2 (1:200 dilution; Lifespan Biosciences, Newark, CA), rabbit anti-S100 (1:100 dilution; AbboMax, Inc., San Jose, CA, USA), anti-rabbit Acetylated Tubulin (1:1000 dilution; Cell Signaling Technology, Danvers, MA, USA) for three overnights at 4°C with gentle agitation. Tissues were rinsed for three changes of 15 minutes with ice-cold 0.25% PBT at 4°C, then incubated in secondary antibody solution consisting of anti-mouse Alexa680 (1:500 dilution; Cell Signaling Technology), anti-rabbit Alexa594 (1:500 dilution; Cell Signaling Technology), and Phalloidin iFluor-488 (1:500 dilution: Abcam, Cambridge, UK) overnight at 4°C with gentle agitation in the dark. The next day, samples were rinsed for three changes of 15 minutes in ice-cold 0.25% PBT at 4°C with gentle agitation under dark conditions.

To label olfactory epithelia with phosphorylated-ERK and total-ERK antibodies (1:500 dilutions each; Cell Signaling Technology, Danvers, MA, USA), antigen retrieval was performed as below, following the initial fixation and 0.25% PBT washes. Tissues were incubated in 150 mM Tris-HCl, pH 9.0 (MilliporeSigma) for 5 minutes at room temperature, then immediately transferred to a 70°C water bath for 15 minutes. Samples were immediately cooled on ice, washed with 0.25% PBT for 15 minutes three times at 4 °C, then incubated in 0.05% Trypsin-EDTA (ThermoFisher, Waltham, MA, USA) for 30 minutes without agitation. Tissues were washed for three changes of 15 minutes with 0.3% PBT at 4°C. The immunohistochemistry protocol then proceeded with the blocking buffer as described above. All experiments used N = 3 biological replicates per antibody combination and morph.

### Activating olfactory sensory neurons by exposing to alanine (pERK detection)

Stone air filters were soaked in fresh, conditioned water for one overnight. The next day, adult *A. mexicanus* surface and Pachón cave morphs were transferred individually to 1 L tanks (24 × 12 × 8 cm polypropylene multi-purpose tanks) and fasted in fresh, conditioned fish water (pH 7.0-7.2, conductivity ∼700 µS) with aeration for 24 hours. The day before the experiment, fish were fasted for 24 hours. L-alanine (MilliporeSigma, concentrations of 10^-10^M, 10^-8^M, or 10^-6^M) or a control (1 μL of 1× PBS) was prepared in the 1L of conditioned fish water (pH 7.0-7.2) in each separate container. After fasting, to remove extra odorants, fish were transferred to clean tanks with fresh conditioned fish water (pH 7.0–7.2) and acclimated for 1.5 hours without aeration. Individual fish were then transferred to the alanine or control containers and allowed to swim freely for 3 minutes (*e.g*., one fish placed in a 10^-6^M L-alanine tank). Then, fish were immediately anesthetized by bathing in ice-cold water for 10 min by monitoring cessation of any gill operculum or body movements. Both left and right sides of olfactory epithelia were quickly dissected and fixed in 4% paraformaldehyde (Electron Microscopy Sciences, Hatfield, PA, USA) in 1× Phosphate-Buffered Saline (Santa Cruz Biotechnology, Inc., Dallas, TX, USA). This procedure was repeated for each treatment concentration and individual fish.

For the time series, the experimental procedures were the same, except for the exposure duration (0, 3, 5, and 10 minutes), which was extended from only 3 minutes, and the use of a single 10 M concentration of L-alanine. The 0-minute alanine treatment served as a baseline control. N = 3 biological replicates per treatment group and morph were used.

### Hybridization Chain Reaction-Fluorescent *in situ* Hybridization (HCR-FISH)

Olfactory epithelia from both Pachón cave and surface morphs of *A. mexicanus* were dissected and fixed in fresh 4% PFA in 1× PBS for 18 hours at 4°C with gentle agitation. The HCR-FISH procedure, including dehydration, rehydration, hybridization, and amplification steps, was adapted from the STAR protocol by Ibarra-García-Padilla et al.^101^. Custom HCR probes targeting *ompa*, *trpc2b*, *v2r*, *taar class1* and *class 3* gene products (mRNA) were designed based on the database sequence (*ompa* and *trpc2b*: XM_022665989.2 and XM_007254133.4, respectively) or sequences detected from our algorithm (*v2r*, *taar class1* and *class 3*: Data S2 and Data S3), and synthesized by Molecular Instruments, Inc. (Los Angeles, CA, USA). Signal amplifications were performed using hairpin amplifiers conjugated to fluorophores: B1-Alexa647, B2-Alexa546, B4-Alexa594, B5-Alexa514, and B5-Alexa488 (Molecular Instruments). All experiments used N = 3 biological replicates per morph.

### Mounting and Sectioning tissues

Following IHC or HCR-FISH, several single lamellae per olfactory epithelial rosette were dissected, and placed on within a circle drawn with PAP pen (ImmEdge Hydrophobic Barrier Pen: Vector Laboratories, Newark, CA, USA) at a slide glass (Corning Incorporated, Corning, NY, USA) using VECTASHIELD PLUS Antifade Mounting Medium (Vector Laboratories, Inc., Newark, CA, USA) and sealed with a cover glass (Fisher Scientific, Pittsburgh, PA, USA).

For cryosections, labeled olfactory epithelial rosettes were immersed in a series of sucrose gradients (VWR International) in 1× PBS: 5% Sucrose in PBS for 3 hours at 4°C, 10% sucrose in PBS for 4 hours at 4°C, and 20% sucrose in PBS overnight at 4°C. Rosettes were embedded in Tissue-Tek O.C.T. Compound (Sakura Finetek USA, Inc., Torrance, CA, USA) and sectioned into 15µm thickness using a Leica CM1950 cryostat (Leica Microsystems, Wetzlar, Germany). Sections were collected on positively charged glass slides (MAS-GP slides: Matsunami Glass Ind., Ltd., Osaka, Japan), dried, and stored at -80 °C until mounted. The sections were rinsed briefly in PBS for 1 minute, and mounted with VECTASHIELD PLUS Antifade Mounting Medium (Vector Laboratories, Inc., Newark, CA, USA) on a slide glass (Corning), sealed with a cover glass (Fisher Scientific). The mounted tissue slides were stored at 4°C protected from light until scanning.

### Quantitative reverse transcription polymerase chain reaction (RT-qPCR)

Adult *A. mexicanus* (surface fish and Pachón cavefish; 4-7 cm in the standard length) were anesthetized in the ice-cold water for at least 7 min, and their both left- and right side olfactory epithelial rosettes were dissected with RNase AWAY-treated dissection tools (ThermoFisher Scientific). The tissue around rosettes was carefully removed under the ice-cold RNase-free PBS (diethyl pyrocarbonate-treated PBS; ThermoFisher Scientific), washed once in the fresh ice-cold RNase-free PBS, recovered in the pre-weighed 1.5 mL tube (SP Bel-Art, Wayne, NJ, USA), and flash frozen by submerging it in liquid nitrogen. All procedures are under RNase-free conditions and both sides of the olfactory epithelia were pooled into one (*i.e*., one tube contains one individual olfactory epithelia). The weight of olfactory epithelia was calculated by subtracting the empty weight of the 1.5 mL tube from the weight of the 1.5 mL tube + tissue. Tissues were stored at -80 °C until use.

Total RNA of olfactory epitheria was extracted via the *Quick*-RNA™ Miniprep Kit (Zymo Research, Irvine, CA, USA). The olfactory epithelia (two rosettes from one individual) were ground by the disposable pestle with a homogenizer (SP Bel-Art) under 300 µL of RNA Lysis Buffer at room temperature. After grinding thoroughly, add 300 µL of fresh RNA Lysis Buffer to bring the total volume to 600 µL. The total RNA was then extracted by following the manufacturer’s protocol.

The mass and quality of extracted total RNA were measured by NanoDrop (Thermo Scientific), and the standardized amount of total RNA was used to reverse-transcribe into cDNA via the iScript™ cDNA Synthesis Kit (Bio-Rad Laboratories, Hercules, CA, USA) according to the manufacturer’s protocol. The synthesized amount of cDNA was also checked by NanoDrop. Quantitative PCR (qPCR) was performed on a CFX96 System (Bio-Rad Laboratories) by using the following method: Each 10uL reaction mix consisted of SsoAdvanced™ Universal SYBR® Green Supermix (Bio-Rad Laboratories), 500 ng of cDNA template, and 500 nM of each forward and reverse primer. Primers were designed using PerlPrimer v1.1.21 with the parameter sets as Tm: 58-62, Amplicon size: 100-300 bases, and checked “GC clamp” and “Exclude %GC” in “Real-time PCR” tab. The primer sequences are available in Data S4. Relative gene expression levels were quantified via the 2^-ΔΔCt^ method and normalized to a panel of housekeeping genes: *b2m*, *eef2a.1*, and *rps18* (Data S4). A melt curve analysis was performed at the end of each run to verify the singularity of the PCR product. All experiments contain three experimental replicates in each of N = 2-3 biological replicates per morph.

### Imaging and Quantitative Analysis

The whole-mounted and cryostat-sectioned samples of immunohistochemistry (IHC) and hybridization chain reaction-fluorescent *in situ* hybridization (HCR-FISH) were imaged using a Leica SP8 X Confocal Laser Scanning Microscope (Leica Microsystems, Wetzlar, Germany). Whole-mounted lamella were scanned from the apical surface to the basal lamina as z-stacks with tiled acquisitions, which were subsequently merged using the LAS X software platform (Leica Microsystems, Wetzlar, Germany). Images were acquired using either a 40×/1.30 oil-immersion objective (whole mount) or a 10×/0.30 dry objective (cryostat sections). For HCR-FISH, acquisition settings (laser power, gain, pinhole) were optimized for each fluorophore combination and maintained constant to enable quantitative comparisons. For IHC samples used for sensory cell counting, acquisition settings were optimized per sample. For pERK signal intensity quantification, samples were imaged with optimized settings, and a mathematical correction factor was applied post-acquisition to normalize for gain differences, as the HyD detector signal response is linear with smart gain. For the pERK time-course series, to reduce scan time, four representative regions of the lamellae with 3 replicates per region (total 12 regions per lamella per fish) (tip, middle, inner zone (R3), and outer zone (R2)) were imaged as single tiles (512x512 pixels) instead of merged tiled lamellae.

### Quantifying Sensory Cells

IHC signals for G_αolf_, G_αo_, S100, and TRPC2 were quantified using ilastik (v1.4.0, Anna Kreshuk’s Lab at the EMBL, Heidelberg, Germany) and Fiji (ImageJ v1.54p, National Institutes of Health). Z-stack projections (maximum intensity) were generated from apical slices with optimal signal intensity using Fiji and imported into ilastik for machine learning-based pixel classification. Each antibody marker was trained separately. G_αolf_ and G_αo_ classifiers were trained to recognize small circular puncta (*i.e.,* dendrites of ciliated and microvillus OSN, respectively), while S100 was trained to identify larger globular cell bodies (*i.e.,* crypt cell bodies). TRPC2-positive cells were classified using three separate classifiers to distinguish between: 1. Punctate TRPC2 signals (dendrites of microvillus OSN); 2. Strong TRPC2-positive cells; 3. Weak TRPC2-positive cells. Strong versus weak TRPC2 signal intensity was determined by visual assessment. Probability maps generated by ilastik were exported and processed in Fiji. Images were converted to 8-bit format, smoothed with a Gaussian blur (σ = 2.0), and despeckled to remove noise. Auto-thresholding (v1.18.0, Intermodes method) followed by watershed segmentation was applied to separate connected objects.

The Particle Analysis tool was used to quantify cell counts with size filters (pixel^2) determined from particle size histograms. Cutoff points according to particle size histograms were estimated based on signal morphology. For TRPC2, additional size thresholds were implemented to distinguish between punctate and larger signals, with size parameters validated for consistency across all samples. ROIs were manually drawn around each whole-mount lamella and further subdivided into three anatomical regions: R1 (outer-most), R2 (outer), and R3 (inner) (*c.f.,* Figure 4). Cell density was calculated as cells per unit area for each ROI.

### Quantifying HCR-FISH Signals

HCR-FISH signals were quantified using Imaris software (v10.2.0, Bitplane AG, Oxford Instruments, Zurich, Switzerland). Confocal z-stacks were imported into Imaris after creating a .ims file in Imaris File Converter and stitching 3D tiles in Imaris Sticher. A surface was generated encompassing the entire stack, and a 3×3×1 median filter was applied to all probe channels: V2R (background subtraction of 7µm), Class 1 TAAR (background subtraction of 3µm), and Class 3 TAARs (background subtraction of 1µm). Another ROI surface was created to exclude non-lamella structures, tissue blemishes, or imaging artifacts. Imaris’ machine learning object classifier was trained to identify signals for each probe. Classifier parameters, including maximum intensity thresholds based on background histogram levels, seed point diameter for estimated cell size, and object volume filters, were optimized for each probe based on signal morphology. The *ompa* signal (green, Figure 6) served as a reference marker for normalization to standardize signal intensity across samples. In detail: 1. Sum intensity of the *ompa* channel was divided by the ROI area to calculate *ompa* sum intensity per unit area; 2. The value of *ompa* sum intensity per unit area was used to normalize the mean and sum intensity statistics of individual detected cells. Absolute signal counts, signal density, and intensity (mean and sum) were quantified for each gene. After these procedures, we found that signals for TAAR3.1, TAAR3.2, and TAAR1 were too weak to detect using the above method (they were buried in background noise). Thus, we summed the signal intensity of each of these genes per lammella, calculated the standardized sum intensity per unit area, and then, standardized with the *ompa* intensity (Figure 6F).

### Quantifying pERK-Positive Cells

pERK-positive cells in both concentration-response and time-course experiments were quantified using Imaris software with a similar process to *Quantifying HCR-FISH Signals*. A surface was generated encompassing the entire stack. Images were processed using the red channel (pERK) with a background subtraction of 2.00 um and a 3×3×1 median filter. To establish a conservative baseline signal intensity, a region of interest (ROI) surface was created using the apical half of the z-stack, excluding non-lamella structures, tissue blemishes, or imaging artifacts. The mean intensity value from this ROI was recorded and applied to the display adjustment window to aid in identifying pERK-positive cells. pERK-positive cells were quantified using the spots classifier tool with the following parameters: estimated XY diameter = 2.00 µm, background subtraction enabled, and z-position filter to retain apical signals. Five representative pERK-positive cells were manually identified, and their mean intensity statistic were used to establish an initial intensity threshold filter. Cells whose intensities were lower than this intensity threshold were excluded from the analysis. Detected pERK-positive cells were manually evaluated to be retained or excluded by referencing the mean intensity statistic of the five reference cells as the minimum intensity range, and the maximum intensity statistic of the detected pERK-positive cells as the maximum intensity range. This classification method was applied consistently across all pERK signal evaluations. All pERK intensity values of pERK-positive cells were standardized with the gain values of its channel. Standardizing pERK intensity with tERK intensity was unnecessary because of the use of the same tissue (olfactory lamella), and rather provided extra noises while quantifying fine intensity shifts of pERK. Absolute cell counts, cell density (cells per unit area), and sum intensity statistics per cell were quantified.

### Particle Tracking

*A. mexicanus* were anesthetized in an ice-cold conditioned fish water, and then, positioned in a custom 3D-printed rectangular chamber with the olfactory pit submerged in the conditioned fish water. The 3D-printed chamber was printed with Adventurer 5M (Zhejiang Flashforge 3D Technology Co., Ltd., Guangdong Province, China). The olfactory pits were imaged in a lateral view (Figure 8) or a dorsal view (Figure 9). The 2.00µm Fluoresbrite® YG Carboxylate Microspheres (Polysciences, Inc., Warrington, PA, USA) were applied in the vicinity of the anterior olfactory pit to visualize water flow patterns toward it. High-speed fluorescent imaging was performed on an BX61WI microscope (Olympus Corp., Tokyo, Japan) equipped with a Hamamatsu ORCA-flash4.0 digital camera (Hamamatsu Photonics K.K., Hamamatsu, Japan) using a 5×/0.15 NA Plan-Fluor objective lens. Images were acquired using MetaMorph Basic software (v7.8.2.0, Molecular Devices, LLC., San Jose, CA, USA). For descriptive flow recordings (Figure 7), high-speed images were recorded at 66.7 frames per s (fps) with a 15ms exposure time for 22.5 s in total. For ciliobrevin-D treatment experiments (Figure 9), images were recorded at 200 fps with a 5 ms exposure time for 5 s. Image scales were calibrated to 2.582µm/pixel.

Particle tracking was performed using the TrackMate plugin (v7.14.0) in Fiji (ImageJ v1.54p, National Institutes of Health) on separate regions of interest (ROIs) drawn for particle localization at the anterior and posterior olfactory pits. Particle detection was performed using the DoG (Difference of Gaussian) detector with a median filter and sub-pixel localization. Tracks were generated using the Advanced Kalman tracker with a maximum gap of 0 frames. For descriptive flow recordings (Figure 7), particle detection used an estimated particle diameter of 7 pixels and a quality threshold of 5. Tracks were generated with an initial search radius and search radius of 5 pixels. Tracks were filtered for duration (below 599.57) and number of spots on track (above 20.55). For Ciliobrevin-D experiments (Figure 9), the detector was set with an estimated particle diameter of 2 pixels and a quality threshold of 50. Tracks were generated with an initial search radius and search radius of 2 pixels. Tracks were filtered for duration (above 31.34) and number of spots on track (above 49.20). Particle velocities were calculated from Max Track Speeds and converted from µm/frame to µm/s using the acquisition frame rate (66.7fps for Figure 7; 200 fps for Figure 9). A custom-made Python script was used to implement a low-pass filter to remove high-frequency whole-image motion (according to the fish motion). All experiments used N = 3 biological replicates per morph and/or treatment group.

### Ciliobrevin-D (dynein inhibitor) Treatment to the Olfactory Pits

40 µM of Ciliobrevin-D in 2% agarose gel was prepared as in follows: 8 µL of the Ciliobrevin stock solution—5 µmol/ mL Ciliobrevin-D (Cytoplasmic dynein inhibitor; MilliporeSigma) in Dimethyl sulfoxide (DMSO; MilliporeSigma) stored in -80 °C and used within one month—were suspended in 1 mL of 0.025% Phenol Red (Fisher Scientific) and 2% UltraPure™ Agarose (Thermo Fisher Scientific) in distilled water at 50 °C. After vortexing, the Ciliobrevin agarose solution was poured into a well of a 24-well dish (Corning) and was stored on ice by sealing it with the plate cover until use. The control agarose gel was prepared by using 8 µL DMSO without Ciliobrevin (1 mL of 0.025% Phenol Red and 2% Agarose). These Ciliobrevin agarose or control agarose are loaded into P10 pipet tip (SureOne™ Micropoint Pipette Tips, Universal Fit; Thermo Fisher Scientific) by jamming it into the agarose gel.

Adult female fish (> 2 years old) were anesthetized in the ice-cold water for at least 5 min and were carefully observed with their motion of gill opercula and fins. After ceasing these motions, the Ciliobrevin or control agarose gel was applied into the olfactory pits from the posterior pits and was loaded until the gel filled the olfactory pits (Figure S8). Olfactory pits were then covered by the Gorilla Super Glue Gel (Gorilla Glue Company, Cincinnati, OH, USA). The agarose gel were treated both sides (behavior assay) or one side for Ciliobrevin agarose and another side for control agarose (flow measurements) per individual fish (Figure S8). Treated fish were recovered in the fresh fish-conditioned water at 22 °C for approximately 30 min.

During the recovery, the glue cap will fell off, and then fish were transferred to the behavior recording stage (behavior assay, See “Behavior assay and analysis in response to alanine”) or reanesthetized with the ice-cold water for the flow imaging (flow measurements). All flow-imaging experiments used N = 3 biological replicates per treatment group and morph.

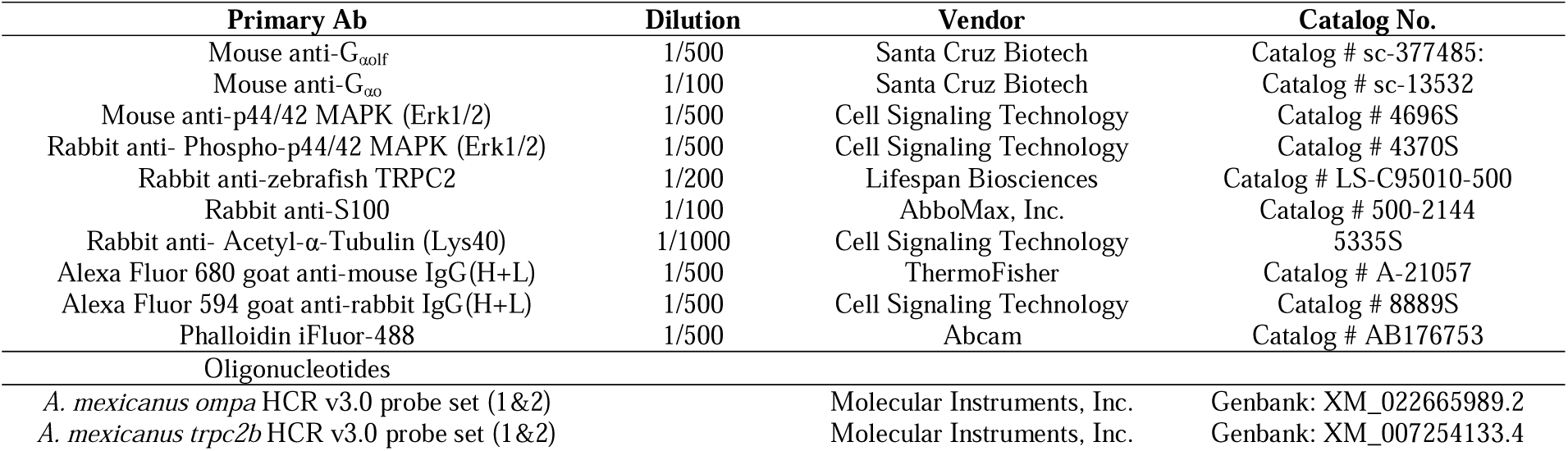
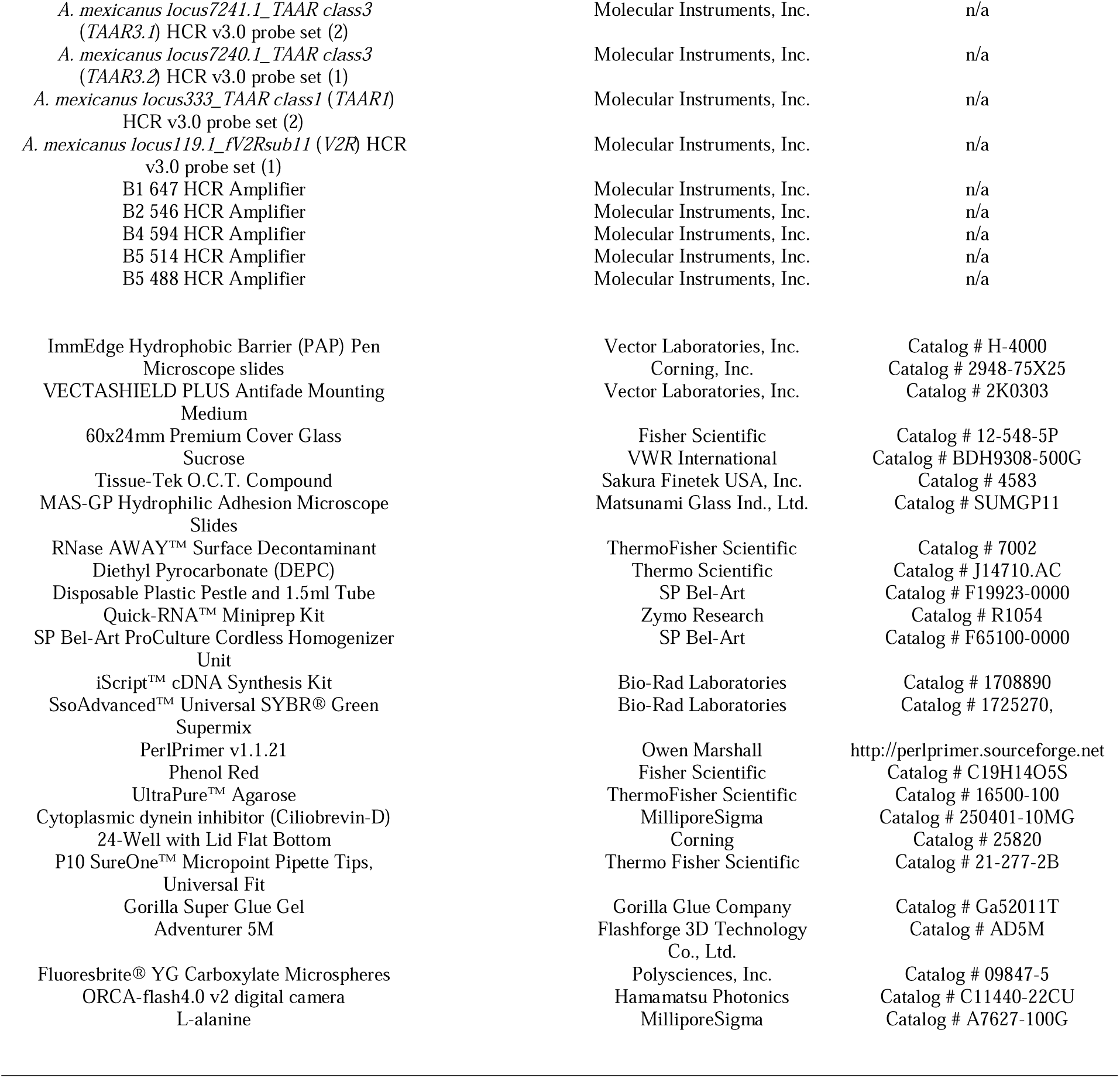

## Acknowledgement

We are grateful to A Jee, B Weise, B Espejo, C Tong, D Inouye, D Burbano, E Stevenson, H Yoshizawa, H Hady, I Matsuyoshi, I Riethbrock, J Southworth, J Boggs, J Cashon, J Hwang, J Tong, K Politan, K Cousens, K Fukumoto, K Wong, M Rai, M Hasegawa, M Quadri, M Pugh, R Chung, S Singh, S Blattner, T Okamura, T Ohta, V Garcia-Tobar, X Barragan Gonzares, X Lolli, and Y Sung for fish care assistance. We also thank to R Peres-David, B Yamamoto, AK Maunakea for their help in PCR and other equipment use, M Norris and T Au la for their qPCR machine use. Lastly, we thank to T Carvalho, K Ewell and O Rivers for the guidance/support for the use of SEM, TEM and a confocal microscope.

## Author information

### Authors and Affiliations

School of Life Sciences, University of Hawai‘i at Mānoa, Honolulu, Hawai‘i, USA

Natalie Choi and Masato Yoshizawa

Bond Life Sciences Center, Division of Animal Sciences, University of Missouri, Columbia, Missouri, USA

Edward S. Ricemeyer and Wesley C. Warren

Chair of Animal Systems Genomics, Faculty of Veterinary Medicine, Ludwig-Maximilians-Universität, Munich, Germany

Edward S. Ricemeyer

School of Natural Sciences, Macquarie University, Sydney, Australia

Maggs X

Institute for the Advanced Study of Human Biology (WPI-ASHBi), Kyoto University, Kyoto, Japan

Zicong Zhang

School of Life Science and Technology, Institute of Science Tokyo, Tokyo, Japan

Masato Nikaido

Department of Surgery, Institute of Data Science and Informatics, University of Missouri, Columbia, Missouri, USA

Wesley C. Warren

### Contributions

NC: designed the experiments, performed the experiment and analyses, and edited the manuscript.

ESR: performed snRNA-seq analysis, edited the manuscript

MX: designed the experiments, performed snRNA-seq experiments and analysis, and edited the manuscript

ZZ: Performed phylogenetic analysis of chemoreceptor genes, edited the manuscript

MN: Performed phylogenetic analysis of chemoreceptor genes, and edited the manuscript

WW: designed the experiments, performed analysis, and edited the manuscript

MY: designed the experiments, performed the experiment and analyses, wrote the initial draft, and edited the manuscript

### Corresponding author

Correspondence to Masato Yoshizawa

### Ethics declarations

Ethics approval and Consent to participate:

The experimental protocols used in this study and fish care were approved by the institutional animal care and use committee (IACUC) at the University of Hawai‘i (17-2560).

Consent to publish:

All authors agreed on publishing these data and this manuscript.

### Author contribution

NC: designed the experiments, performed the experiment and analyses, and edited the manuscript.

ESR: performed snRNA-seq analysis, edited the manuscript

MX: designed the experiments, performed snRNA-seq experiments and analysis, and edited the manuscript

ZZ: Performed phylogenetic analysis of chemoreceptor genes, edited the manuscript

MN: Performed phylogenetic analysis of chemoreceptor genes, and edited the manuscript

WW: designed the experiments, performed analysis, and edited the manuscript

MY: designed the experiments, performed the experiment and analyses, wrote the initial draft, and edited the manuscript

### Data Availability

The video and image datasets, and their analysis scripts generated and/or analyzed during the current study are available at the university’s shared server and will be deposited to Dataverse https://doi.org/10.7910/DVN/DMOOBC. FATE program is available at https://github.com/Hikoyu/FATE. All snRNA-seq reads and count matrices are available on NCBI Gene Expression Omnibus under accession GSE319893. The annotations, multiple sequence alignments, and phylogenetic trees for chemosensory receptors are available on Zenodo (https://doi.org/10.5281/zenodo.18691471).

---

**Figure S1.**
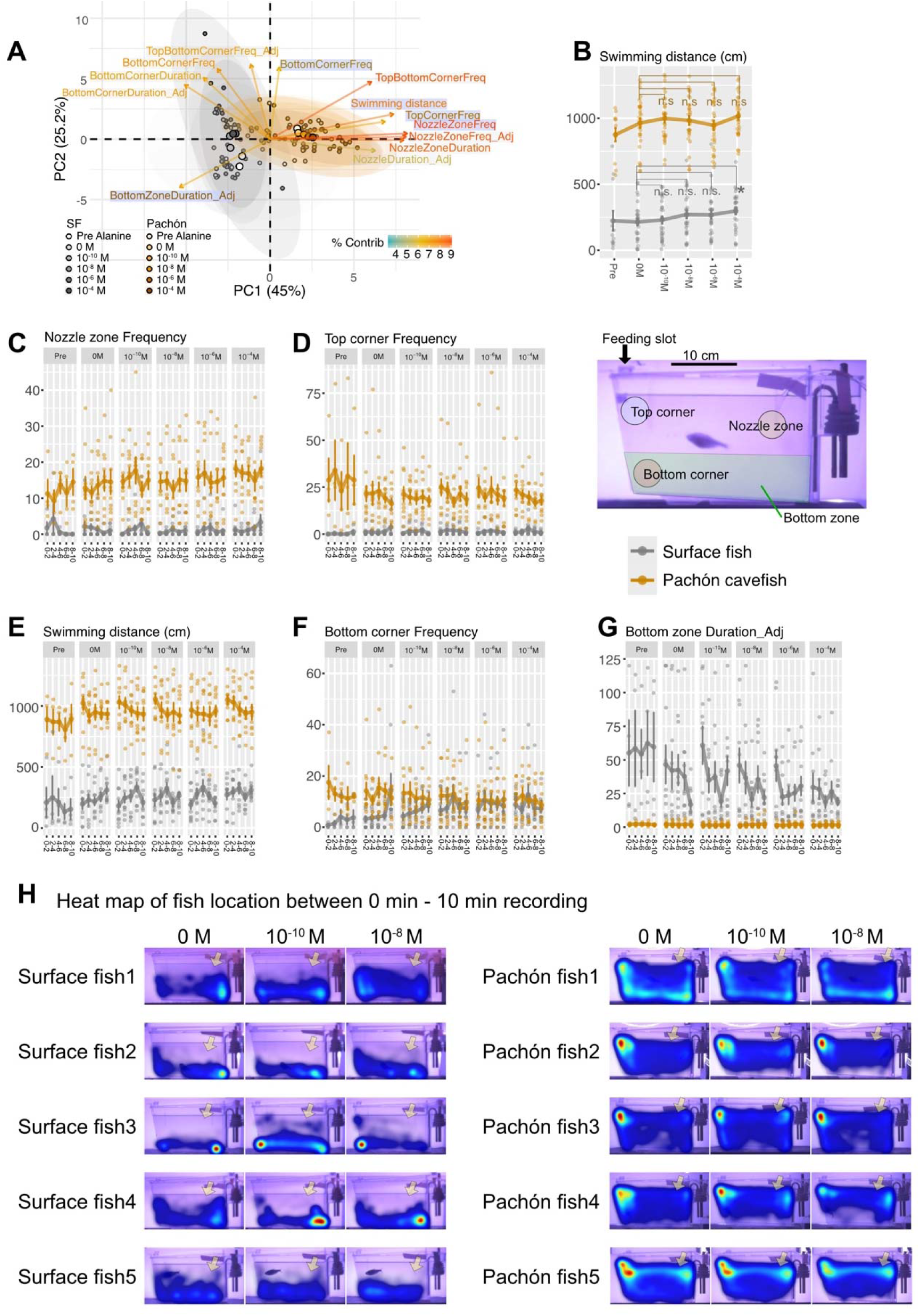
Behavior scores in response to the alanine injection. (A) Principal component analysis (PC1 and PC2) of different behavior scores revealed the responsible loading vectors of behavior parameters that describe the alanine response from zero to 10^-4^ M of alanine. Each fish was exposed to no alanine (‘pre’ treatment experiment, and injection of 3 mL ‘0 M’ alanine [no alanine] solution to the fish tank), followed by incremental concentration of alanine solution every 10 min (10^-10^ M, 10^-8^ M, 10^-6^ M, and 10^-4^ M). Blue banner loading vectors were significant and drove the difference between surface fish and cave fish. Colors of loading vectors represent the percentage contribution to the variation. Behavior parameters were measurements of visiting frequencies and total staying duration in each of these four zones. (B) Swimming distance in the first 6 min. Cavefish did not significantly increase swimming distance while surface fish did at 10^-4^ M alanine injection (X^2^(1) = 8.7, P = 0.01253; Data S6). (C) Time course of visiting frequency to the nozzle zone (alanine source) in the bins of 0-2 min, 2-4 min, 4-6 min, 6-8 min, and 8-10 min. Different injected alanine concentrations (0 M, 10^-10^ M, 10^-8^ M, 10^-6^ M, and 10^-4^ M) and no injection (Pre) were shown. The same data, summing 0-6 min scores, were used in Figure 1C. (D) Time course of visiting frequency to the top corner in the bins of 0-2 min, 2-4 min, 4-6 min, 6-8 min, and 8-10 min. The same data, summing 0-6 min scores, were used in Figure 1D. (E) Time course of swimming distance in the bins of 0-2 min, 2-4 min, 4-6 min, 6-8 min, and 8-10 min. The same data, summing 0-6 min scores, were used in the panel B. (F) Time course of visiting frequency to the bottom corner in the bins of 0-2 min, 2-4 min, 4-6 min, 6-8 min, and 8-10 min. The same data, summing 0-6 min scores, were used in Figure 1F. (G) Time course of staying duration in the bottom zone in the bins of 0-2 min, 2-4 min, 4-6 min, 6-8 min, and 8-10 min. The same data, summing 0-6 min scores, were used in Figure 1E. (H) Examples of 5 fish localizations during 10 min of recording (heatmap).

**Figure S2.**
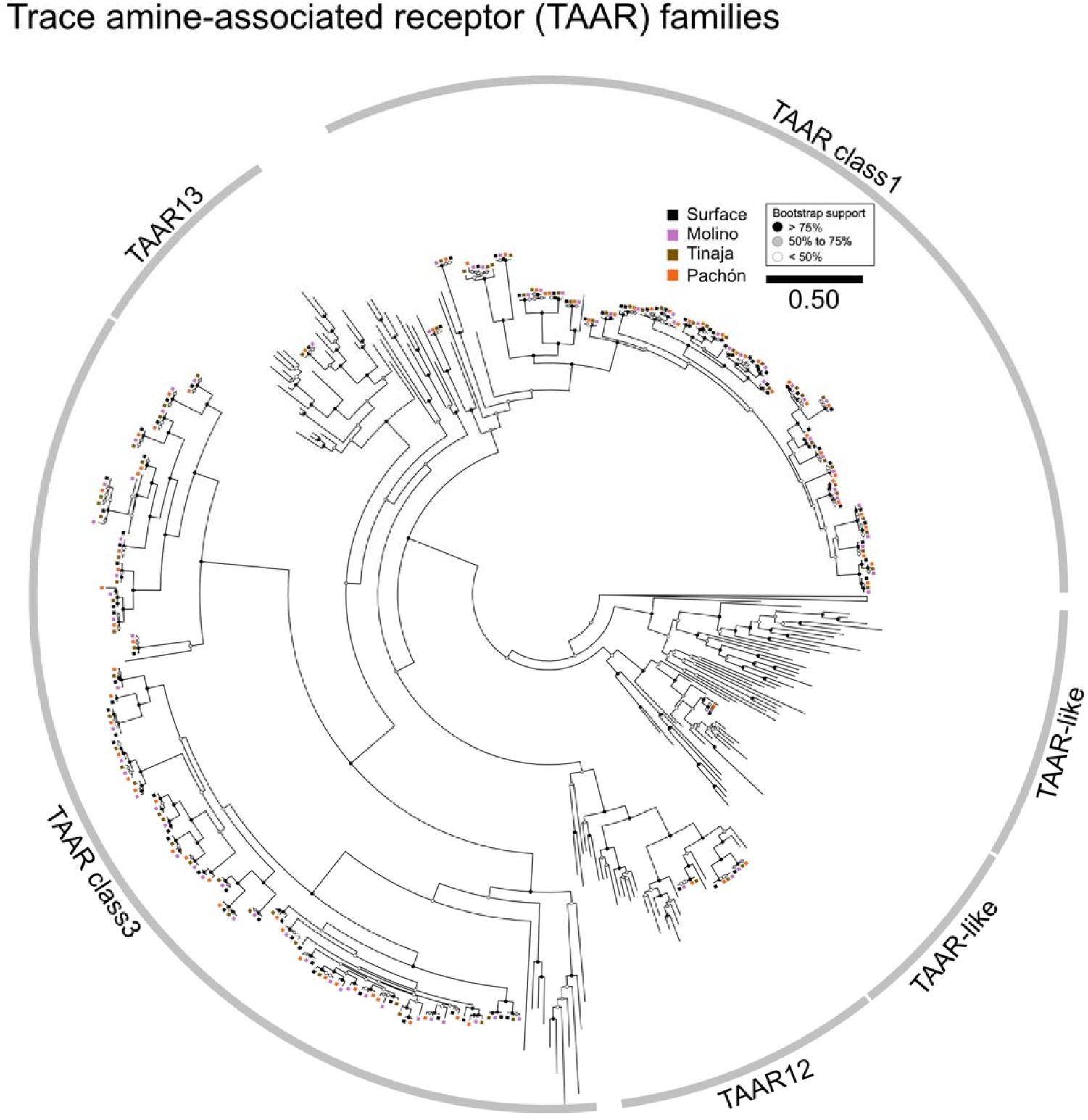
Phylogenetic tree of Trace Amine-Associated Receptors (TAARs) in *Astyanax mexicanus*.

**Figure S3.**
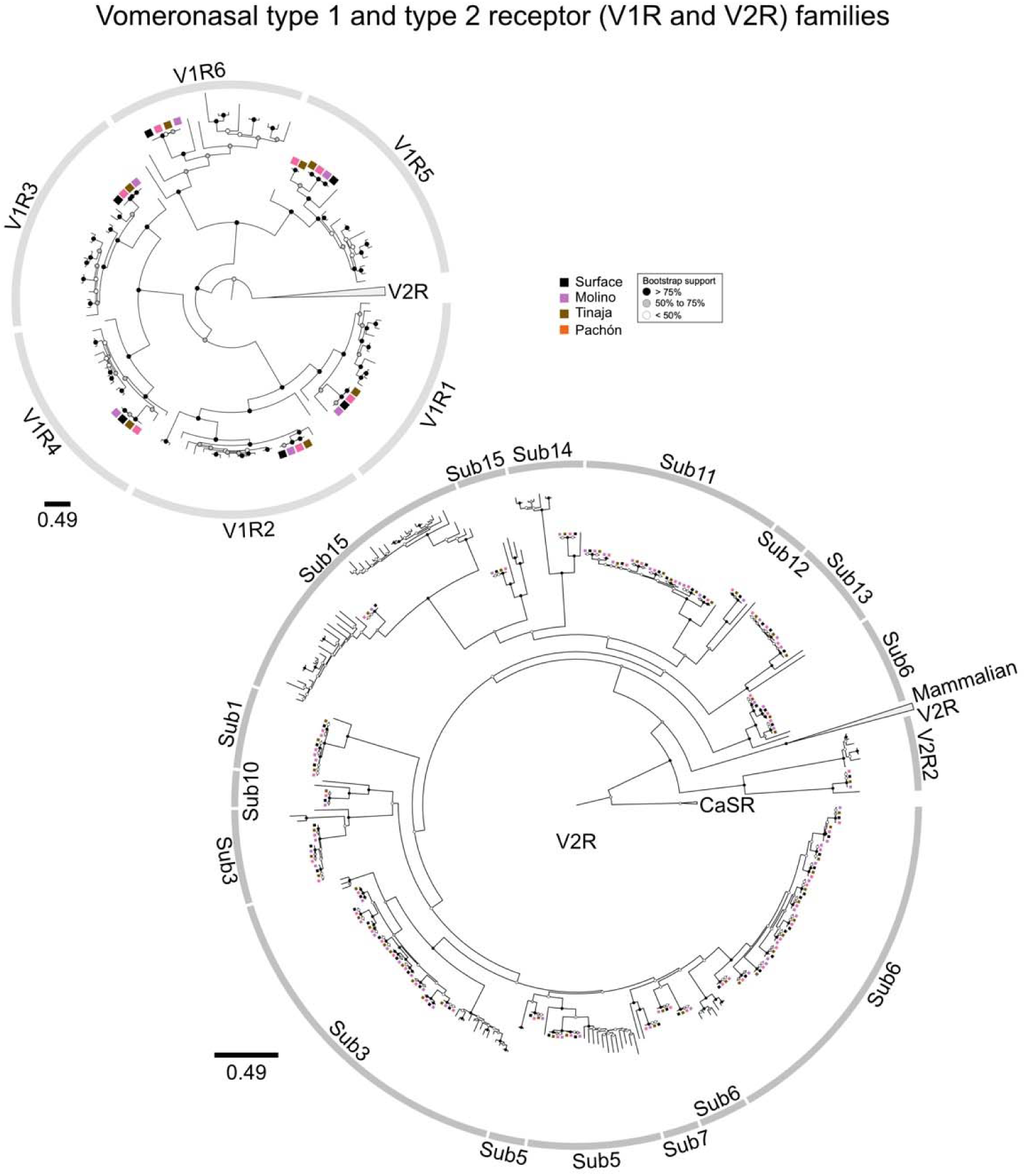
Phylogenetic tree of Vomeronasal receptors, V1R and V2R, in *Astyanax mexicanus*.

**Figure S4.**
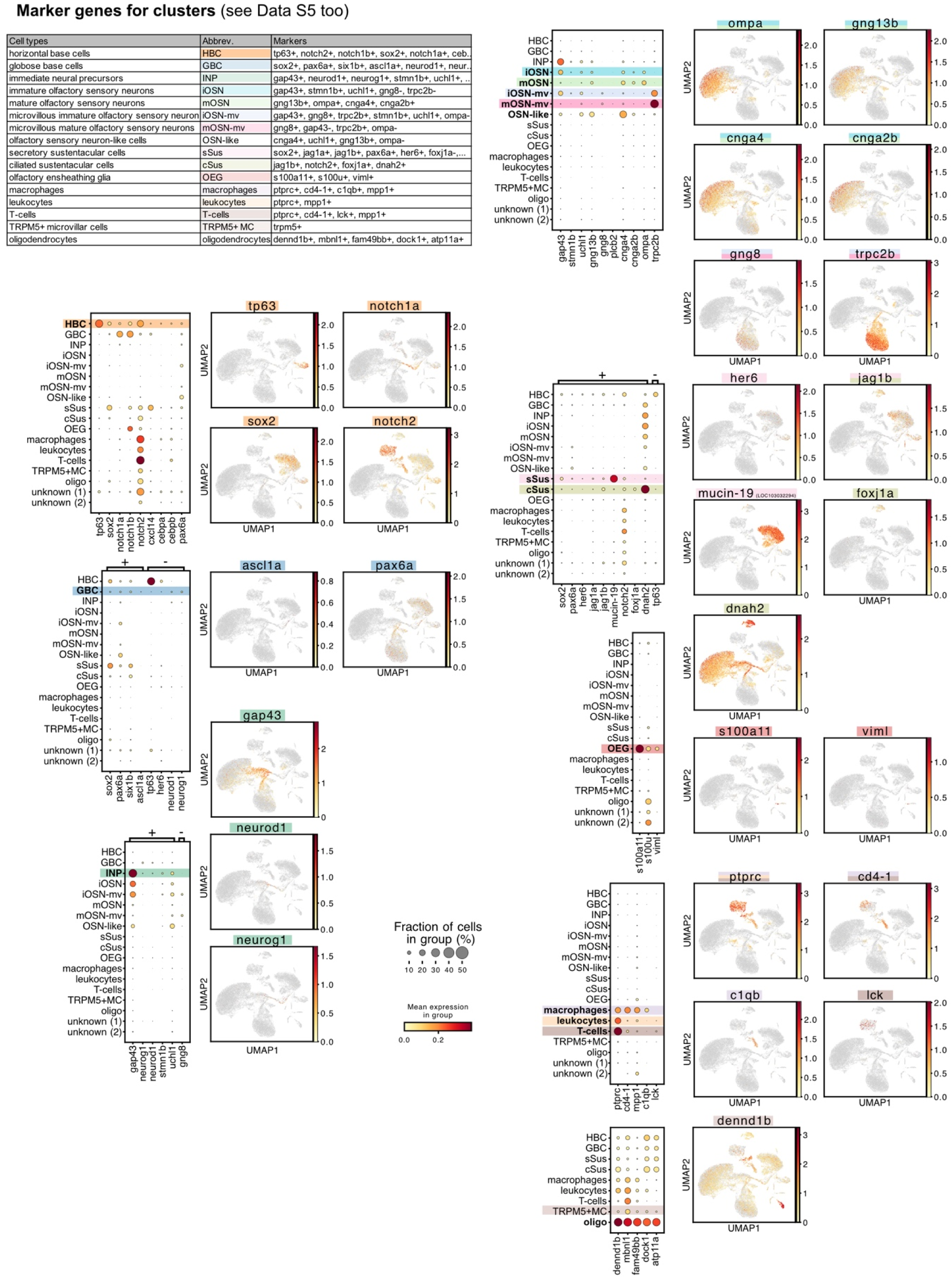
Characterizing each cluster for the result of single-nucleus RNA sequencing. Top left: marker genes for each cluster, characterized in this study. Please also see Data S5. Bubble plots (left panels) and expressed clusters (right panels) for marker genes/ differentially expressed genes compared with other clusters. Conserved marker genes were detected in *A. mexicanus* too (*e.g*., *tp63*, *sox2*, *notch* in HBC, *omp* and *gng13* in mOSN; see Data S5 too). cSus: ciliated sustentacular cells, GBC: Globose basal cell, HBC: horizontal basal cell, INP: immediate neuronal precursor, iOSN: ciliated immature olfactory sensory neuron, iOSN-mv: micrivillus immature olfactory sensory neuron, mOSN: ciliated mature olfactory sensory neuron, mOSN-mv: microvillus mature olfactory sensory neurons, sSus: secretory sustentacular cell, OEG: olfactory ensheathing glia, TRPM5+MC: TRPM5-positive microvillar cell.

**Figure S5.**
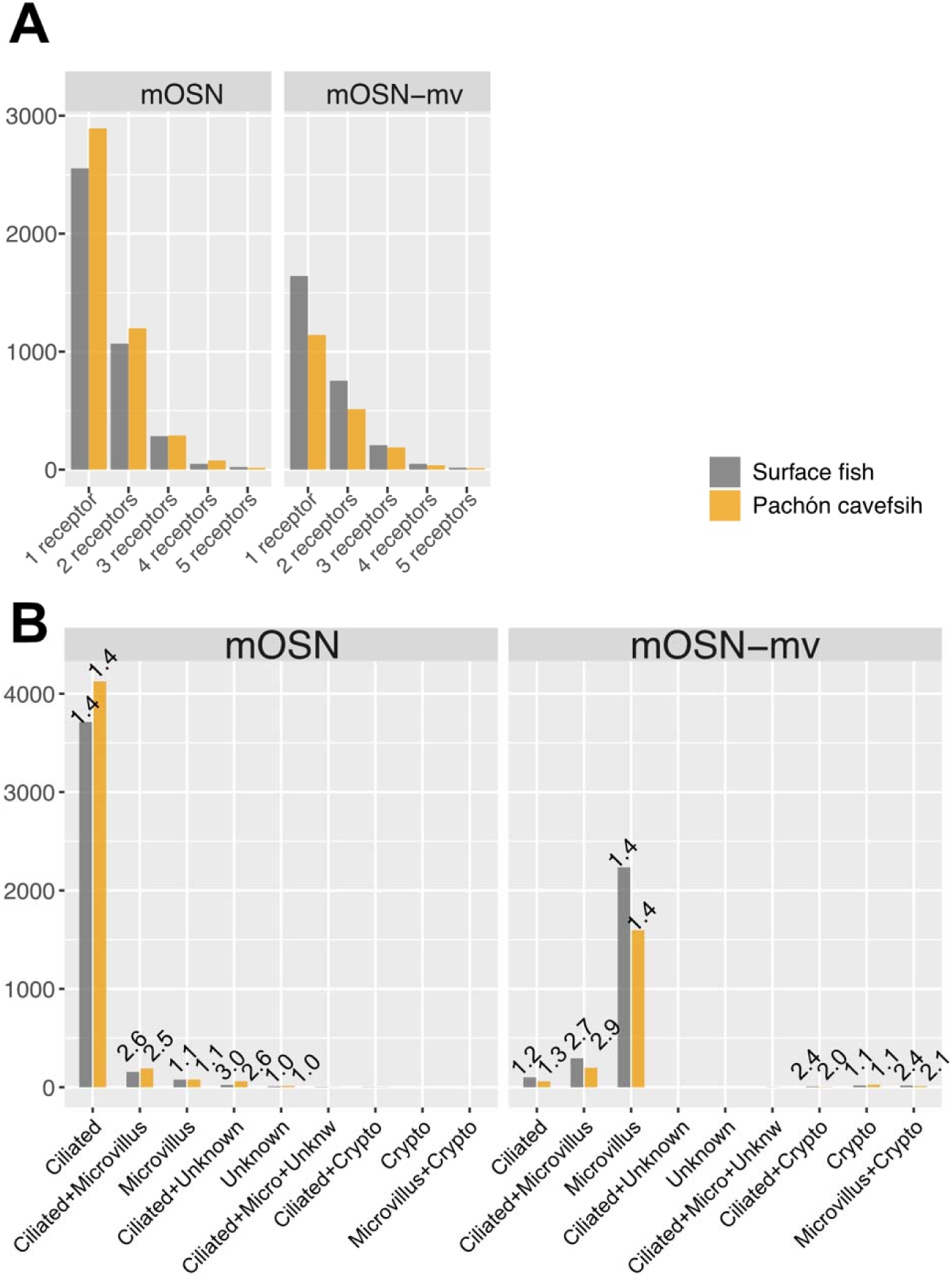
Many of *A. mexicanus* olfactory sensory neurons (OSN) express one receptor gene per OSN according to the result of single-nucleus RNA sequencing. (A) Mature OSNs, ciliated (mOSN, left) and microvillus (mOSN-mv, right), expressed a single receptor gene (1 receptor) or two in surface fish and Pachón cavefish. (B) The chemoreceptor gene numbers per cell were categorized by receptor genes for ciliated (ORs and TAARs), microvillus (V2Rs), crypt (V1Rs), and unknown (T2R). The average numbers of receptor genes expressed in each category per cell are shown at the top of the bar graphs. The cells expressing the single categories of dedicated chemoreceptors (OR and TAAR genes for ciliated mOSN or V2R genes for microvillus mOSN-mv) expressed more than one gene on average (1.4 genes), indicating multiple receptor genes in the same category may be expressed per cell. In contrast, cells expressing genes in the different categories (*e.g*., Ciliated+Microvillus) are in much smaller proportions. No significant difference between surface fish and Pachón cavefish was observed.

**Figure S6.**
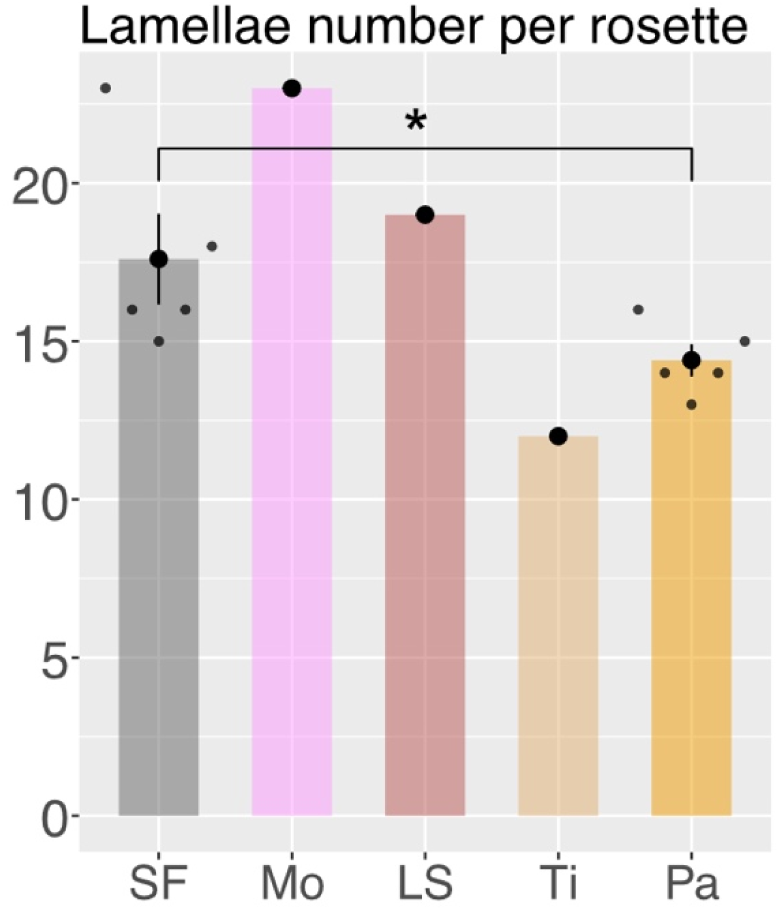
Number of lamellae in the olfactory rosettes of *A. mexicanus* populations. Scanning electron microscope images were used for counting the number of olfactory lamellae. Surface fish (SF) N = 5, Molino cavefish (Mo) N = 1, Sabinos cavefish (LS) N = 1, Tinaja cavefish (Ti) N = 1, and Pachón cavefish (Pa) N = 5 were used. All female individuals. Pachón cavefish lamellae were significantly fewer than those of surface fish (W = 22.5, P-value = 0.03334). *: P < 0.05.

**Figure S7.**
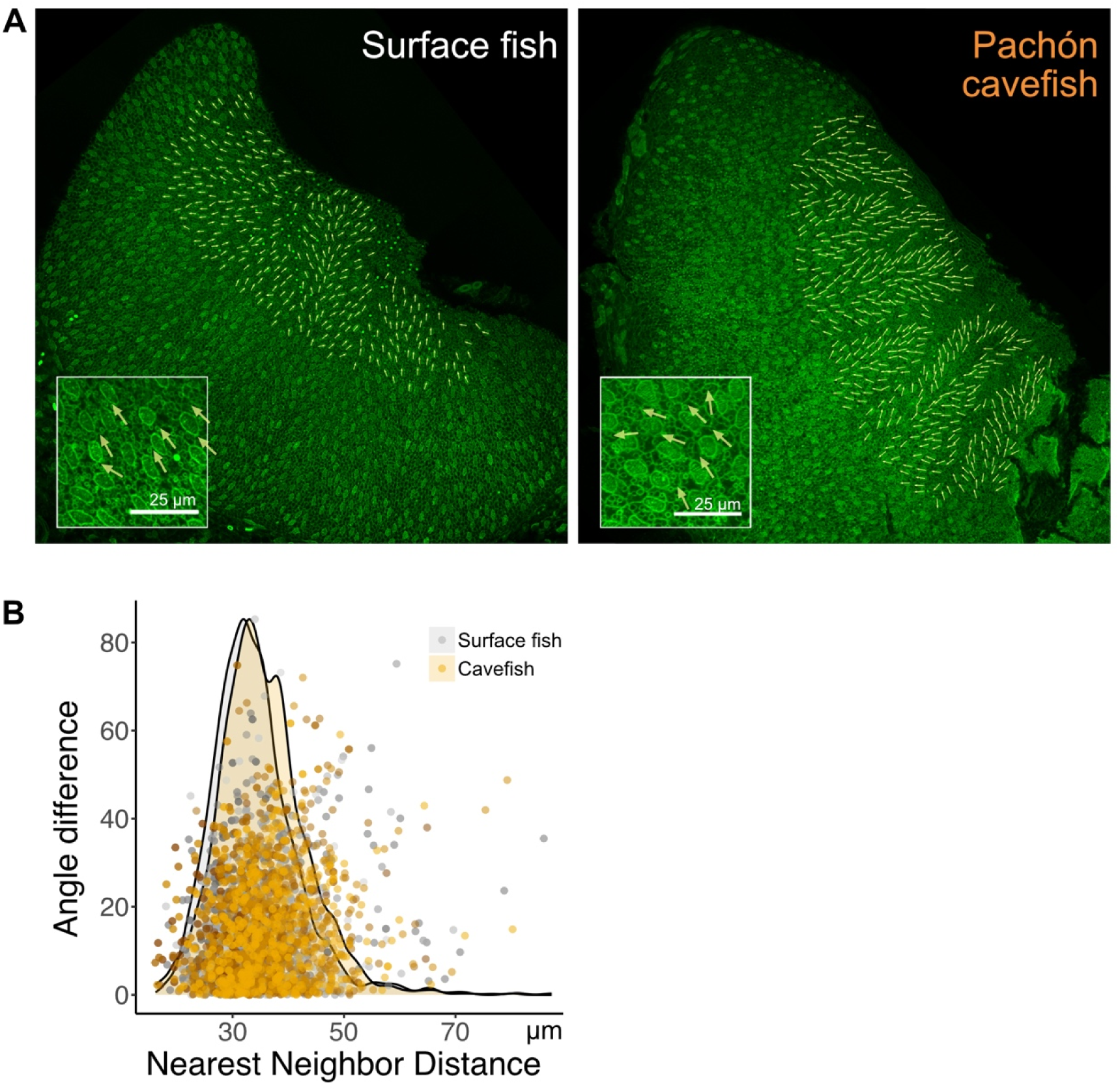
Actin network that likely lines motile cilia implies the direction of the motile cilia stroke. (A) A ladder-like lining of actin network, which is also observed in the tracheal cilia cells^102,103^, indicates the orientation of motile cilia motion (insets). The orientation of motile cilia cells in Pachón cavefish appears disturbed. Lines and arrows indicate the orthogonal axis to the actin ladder network. (B) The angle differences against the nearest neighbor distances between two motile cilia cells were plotted. Cavefish density distribution has a secondary peak, indicating cavefish have a motile cilia cell population that is more apart from each other and at a different angle than surface fish. Surface fish and cavefish distribution was different in their nearest neighbor distance–Angle difference distributions (energy distance test with 999 permutations, E = 181.7, P = 0.001). Further detailed study is needed to confirm the orientations of motile cilia. N = 4 lamellae per population.

**Figure S8.**
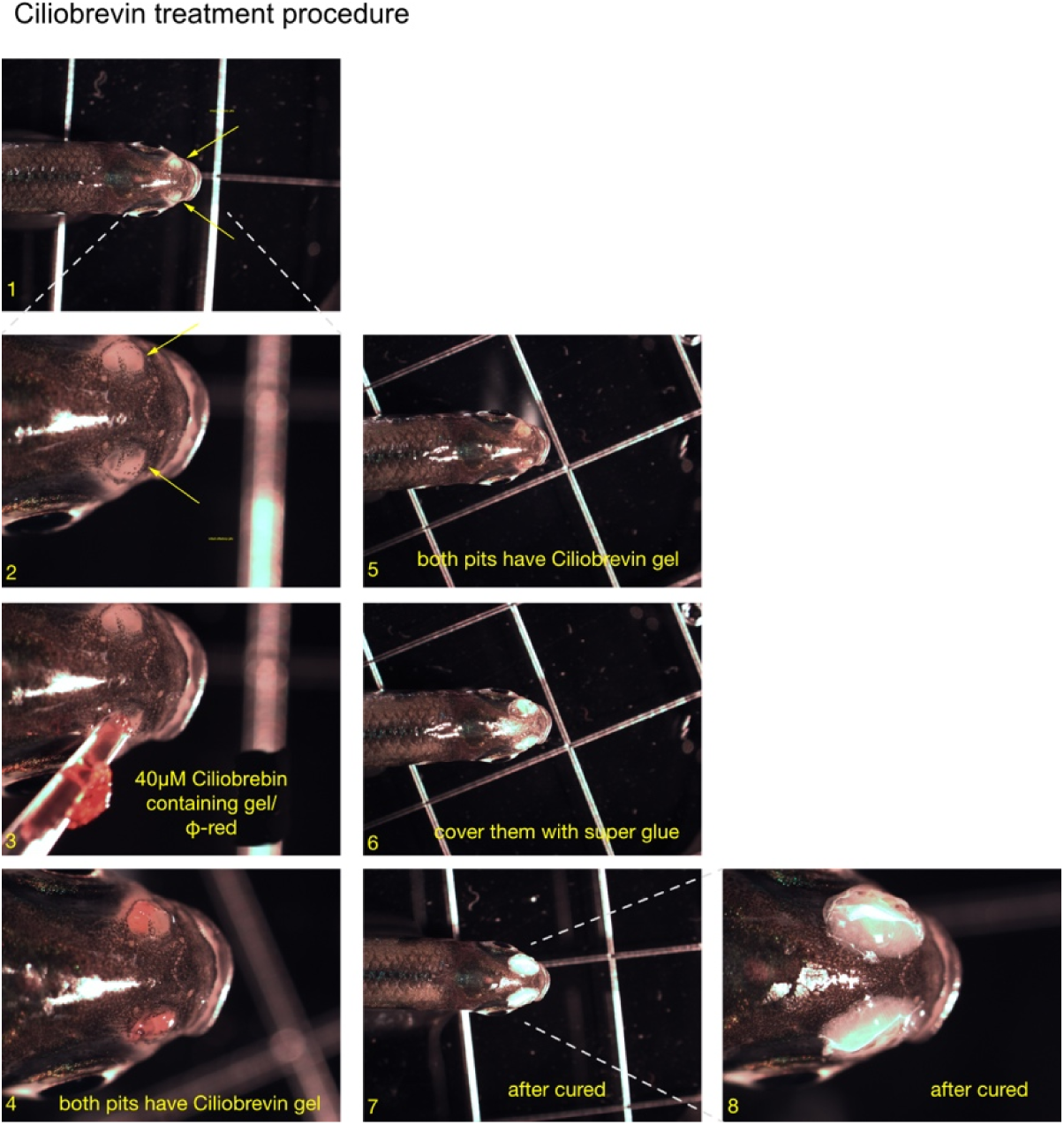
Ciliobrevin treatment at the nasal pits. Treatment procedure for ciliobrevin-D to the olfactory pits. 1. Fish were anesthetized with ice-cold water, and hold it upright position. 2. olfactory pits were indicated with arrows. 3. 40 µM ciliobrevin-D in 2 % agarose with 0.025% phenole red was air-injected into the posterior olfactory pit using a micropipette. 4.-5. After loading the ciliobrevin gel 6. Olfactory pits were covered by superglue 7. After curing (∼1 min), the fish were released in the fresh fish-conditioned water at ∼22°C, and their recovery was carefully observed until they actively swam. The superglue covers eventually fell off ∼30 min after curing.

**Figure S9.**
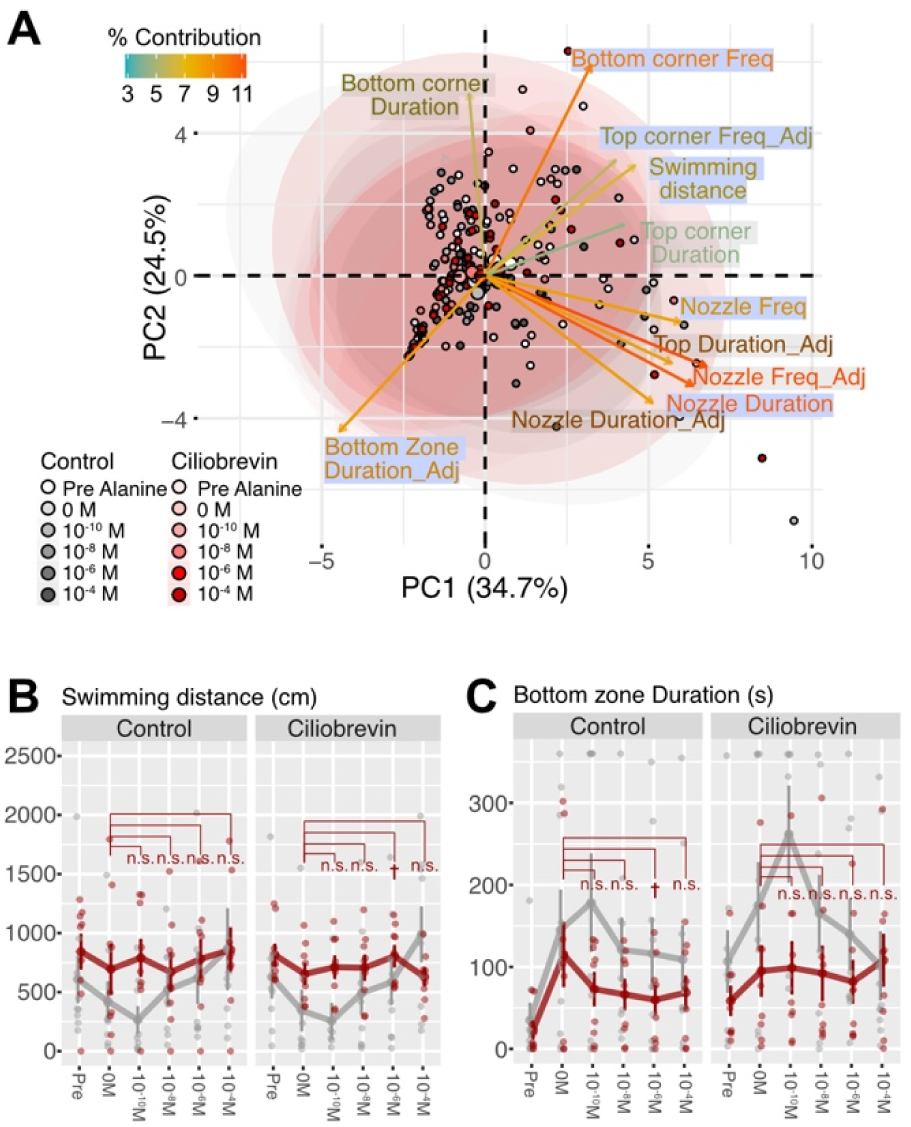
Behavior scores of Ciliobrevin-treated surface fish in response to the alanine injection. (A) PCA plot of control and ciliobrevin-treated surface fish. The top loading vectors are blue bannered: Nozzle zone duration, Bottom corner frequency, and Bottom zone duration. (B) Swimming distance in the first 6 min. Either ciliobrevin-treated or control surface fish increased swimming by the alanine injection (Data S6). (C) Staying duration in the Bottom zone in the first 6 min. No detectable change compared with 0 M alanine in either fish (see Data S6). †: P < 0.05 before the post-hoc correction but P > 0.05 after it, *: P < 0.05, **: P < 0.01, ***: P < 0.001.

**Data S1. Differentially express gene analysis by pooling cell cluster identified in snRNA-seq**

**Data S2. Gene sequences used for designing HCR-FISH probes**

**Data S3. All detected chemoreceptor sequences (transcripts)**

**Data S4. Primer sets used for RT-qPCR**

**Data S5. Identification and characterization of snRNA-seq clusters and their references.**

**Data S6. Details of statistical scores for Figures 1, 2, 4, 6, 8, 9.**

