## Supplemental data S1-S6 for "Novel mechanisms of chemosensory adaptation to the cave environment": Data S5.html

2-annotation


### Introduction¶

In this notebook, I'll be annotating the results of clustering single nuclei
data from the olfactory epithelium of 2 cave and 2 surface samples.

First, a list of cell types we expect to find in this dataset, along with
marker genes and references:

- Horizontal basal cells: TP63+, SOX2+, PAX6+ (reviewed in Schwob et al., 2017)
  - Also NOTCH1+, NOTCH2+
  - Cxcl14+, Cebp+ (Durante 2020)
  - See also Chen et al. (2019)
- Globose basal cells: SOX2+, PAX6+, SIX1+, ASCL1+, TP63-
  - Schwob et al. (2017) break these down into different stages and lineages,
    but there is not consensus about this yet, so perhaps best to just have a
    GBCs category
  - There also may be KIT+ GBCs (Goss et al., 2016)
- Immediate neural precursors (INP)
  - Fletcher et al. (2017) differentiate between early INP (neurod1+,
    neurog1+) and late INP (gng8+)
- Olfactory sensory neurons: UCHL1+, NST+, OR+, OMP+ (Schwob et al., 2017)
  - Immature olfactory sensory neurons (iOSN): GAP43+, GNG8+ (Durante et al.,
    2020)
  - Mature olfactory sensory neurons (mOSN): GNG13+ (Durante et al., 2020)
  - CNG-family genes + in catfish (Goulding et al., 1992)
  - CNGA2+ (Hanchate et al., 2015)
  - TRPC2+ in zebrafish (Calvo-Ochoa et al., 2021)
  - More markers from mouse in Tan et al. (2015)
- Sustentacular cells: SOX2+, PAX6+, HES1+ (Schwob et al., 2017)
  - Also JAG1+, NOTCH2+ (but so do HBCs, so NOTCH1-/TP63- is the key)
  - Triana-Garcia et al. (2021) differentiate between ciliated and secretory
    sustentacular cells in delta smelt
  - Markers for the ciliated ones include DNAH9 and FOXJ1
- Microvillar cells: heterogeneous, reviewed by Bryche et al. (2021)
  - TRPM5+ MCs (Genovese & Tizzano, 2018)
  - PLCB2+ MCs
  - TRPC6+ MCs
  - Calvo-Ochoa et al. (2021) report TRPC2+/OMP- microvillous OSNs in zebrafish
- Ensheathing glia: NTFR+ (also Schwann and fibroblasts), S100+, VIM+, SOX10+
  (Oprych et al., 2017)
  - These probably do exist in fish, although they are poorly characterized
    (Lazzari et al., 2013)
- Multipotent duct/gland cells: PAX6+, SOX9+ (Schwob et al., 2017)
- Epithelial cells (Schwob et al., 2017)
- Various immune cells

Fish *do not* have Bowman's glands (Getchell & Getchell, 1992).

In [1]:

```
from collections import defaultdict
import warnings

import matplotlib
import pandas as pd
import scanpy as sc
from matplotlib.colors import LinearSegmentedColormap

%matplotlib inline
matplotlib.rcParams["figure.figsize"] = (7.0, 7.0)

warnings.filterwarnings('ignore')

cm = LinearSegmentedColormap.from_list("Custom", [(0.9, 0.9, 0.9), (0.7, 0, 0)], N=40)
```

In [2]:

```
markers = pd.read_csv("all_markers.csv")
markers
```

Out[2]:

|  | cluster | gene | pval\_adj | LFC | pct.1 | pct.2 |
| --- | --- | --- | --- | --- | --- | --- |
| 0 | 0 | ebf1a | 0.000000 | 4.170580 | 98.555957 | 34.713633 |
| 1 | 0 | atp2b2 | 0.000000 | 2.822871 | 97.584004 | 55.411401 |
| 2 | 0 | adcy8 | 0.000000 | 4.239890 | 88.281033 | 21.356547 |
| 3 | 0 | ebf1b | 0.000000 | 2.419339 | 96.445432 | 57.229766 |
| 4 | 0 | umodl1 | 0.000000 | 2.702160 | 91.835601 | 47.348077 |
| ... | ... | ... | ... | ... | ... | ... |
| 36746 | 28 | arhgef18a | 0.009823 | 4.489867 | 13.432836 | 1.179753 |
| 36747 | 28 | atp2a2a | 0.009856 | 4.962668 | 13.432836 | 0.816752 |
| 36748 | 28 | pkp3a | 0.009889 | 3.068655 | 14.925373 | 3.045173 |
| 36749 | 28 | arhgap46b | 0.009936 | 2.116555 | 20.895522 | 7.586045 |
| 36750 | 28 | cldn7b | 0.009954 | 1.461350 | 28.358209 | 17.528233 |

36751 rows × 6 columns

In [3]:

```
adata = sc.read_h5ad("clusters_original.h5ad")
```

In [4]:

```
sc.pl.umap(adata, color="leiden", legend_loc="on data")
```

### Horizontal basal cells¶

Horizontal basal cells are:

- TP63+, SOX2+, PAX6+ (Schwob et al., 2017)
- Schwob et al. (2017) also say NOTCH1+ and NOTCH2+
- KRT5+ is used in zebrafish (Kocagöz et al., 2022)
- CXCL14+, CEBP+ (Durante 2020)
- See also Chen et al. (2019)

Some orthology notes:

- only pax6a is orthologous with pax6 in mammals (not pax6b)
- only six1b is orthologous with mammalian six1 (not six1a)
- only ascl1a is orthologous with mammalian ascl1 (not ascl1b)
- notch1a/b are both orthologous with mammalian notch1
- there is no Amex ortholog of zebrafish KRT5

In [5]:

```
sc.pl.dotplot(
    adata,
    ["tp63", "sox2", "notch1a", "notch1b", "notch2", "cxcl14", "cebpa", "cebpb", "pax6a"],
    "leiden",
    swap_axes=True,
)
```

In [6]:

```
markers[
    markers.cluster.isin([17, 22])
    & markers.gene.isin(
        ["tp63", "sox2", "notch1a", "notch1b", "notch2", "cxcl14", "cebpa", "cebpb"]
    )
]
```

Out[6]:

|  | cluster | gene | pval\_adj | LFC | pct.1 | pct.2 |
| --- | --- | --- | --- | --- | --- | --- |
| 23769 | 17 | tp63 | 4.443099e-49 | 7.786191 | 45.010183 | 1.139419 |
| 23942 | 17 | notch2 | 9.151801e-23 | 2.023822 | 36.863544 | 9.296854 |
| 24268 | 17 | notch1b | 2.233764e-11 | 2.379936 | 16.700611 | 3.485480 |
| 24405 | 17 | sox2 | 2.945207e-09 | 2.024155 | 16.293279 | 4.947567 |
| 24635 | 17 | notch1a | 5.227671e-07 | 2.930173 | 8.757637 | 1.199919 |
| 25264 | 17 | cebpa | 1.335059e-03 | 2.802177 | 3.869654 | 0.695752 |
| 29897 | 22 | notch2 | 1.183026e-17 | 2.135093 | 42.307692 | 9.296854 |
| 30446 | 22 | tp63 | 9.037422e-06 | 3.172862 | 9.935897 | 1.139419 |
| 30770 | 22 | cebpa | 2.459337e-04 | 3.578737 | 7.371795 | 0.695752 |

Cluster 17 is + for all HBC markers except cxcl14, so labeling this HBC.

### Globose basal cells¶

Globose basal cells are:

- SOX2+, PAX6+, SIX1+, ASCL1+, TP63-, HES1- (Schwob et al., 2017)
- NEUROD1-, NEUROG1- (Fletcher et al., 2017)

Orthology notes:
Some orthology notes:

- only pax6a is orthologous with pax6 in mammals (not pax6b)
- only six1b is orthologous with mammalian six1 (not six1a)
- only ascl1a is orthologous with mammalian ascl1 (not ascl1b)
- her6 is the ortholog of mammalian HES1

In [7]:

```
sc.pl.dotplot(
    adata,
    {
        "+": ["sox2", "pax6a", "six1b", "ascl1a"],
        "-": ["tp63", "her6", "neurod1", "neurog1"],
    },
    "leiden",
    swap_axes=True,
)
```

In [8]:

```
markers[markers.gene.isin(["sox2", "pax6a", "six1b", "ascl1a"])]
```

Out[8]:

|  | cluster | gene | pval\_adj | LFC | pct.1 | pct.2 |
| --- | --- | --- | --- | --- | --- | --- |
| 574 | 0 | pax6a | 7.203564e-10 | 1.067149 | 4.720911 | 3.290535 |
| 1732 | 2 | sox2 | 2.938943e-114 | 3.372988 | 24.056095 | 4.947567 |
| 1814 | 2 | six1b | 2.193570e-91 | 3.787138 | 18.950018 | 3.478758 |
| 2722 | 2 | pax6a | 3.759023e-21 | 1.711320 | 8.234448 | 3.290535 |
| 10745 | 8 | sox2 | 2.853380e-76 | 3.285533 | 32.565056 | 4.947567 |
| 11143 | 8 | six1b | 3.242661e-32 | 2.708711 | 16.579926 | 3.478758 |
| 11944 | 8 | pax6a | 2.518590e-10 | 1.501439 | 8.921933 | 3.290535 |
| 15544 | 11 | pax6a | 9.688556e-12 | 1.973917 | 12.264151 | 3.290535 |
| 18757 | 13 | six1b | 2.464879e-06 | 1.300520 | 13.518198 | 3.478758 |
| 20190 | 14 | sox2 | 2.878413e-16 | 2.005172 | 26.306620 | 4.947567 |
| 20429 | 14 | six1b | 3.085221e-09 | 1.688426 | 16.550523 | 3.478758 |
| 24405 | 17 | sox2 | 2.945207e-09 | 2.024155 | 16.293279 | 4.947567 |
| 25378 | 17 | six1b | 2.756719e-03 | 1.158631 | 8.350305 | 3.478758 |
| 33661 | 24 | pax6a | 5.892760e-07 | 2.345204 | 18.725100 | 3.290535 |

In [9]:

```
markers[markers.gene.isin(["tp63", "her6", "neurod1", "neurog1"])]
```

Out[9]:

|  | cluster | gene | pval\_adj | LFC | pct.1 | pct.2 |
| --- | --- | --- | --- | --- | --- | --- |
| 3677 | 2 | her6 | 1.004953e-07 | 2.149847 | 2.265372 | 0.742807 |
| 12072 | 8 | her6 | 5.786568e-09 | 2.963772 | 4.163569 | 0.742807 |
| 14584 | 10 | neurod1 | 1.188067e-07 | 4.744189 | 3.359173 | 0.289056 |
| 14836 | 10 | neurog1 | 1.554705e-04 | 3.349569 | 2.153316 | 0.299140 |
| 22364 | 16 | neurog1 | 6.419275e-07 | 5.586073 | 7.171315 | 0.299140 |
| 22617 | 16 | neurod1 | 2.957134e-05 | 4.264730 | 4.980080 | 0.289056 |
| 23769 | 17 | tp63 | 4.443099e-49 | 7.786191 | 45.010183 | 1.139419 |
| 24404 | 17 | her6 | 2.859727e-09 | 4.105448 | 10.183299 | 0.742807 |
| 30446 | 22 | tp63 | 9.037422e-06 | 3.172862 | 9.935897 | 1.139419 |

16 is the closest to the definition of GBC, but it is also NEUROD1+/NEUROG1+,
which are markers for differentiating but still proliferating cells. The best
explanation we can come up with for this is that fish cells may retain
proliferation capacity later than mice. So calling 15 GBCs.

### Immediate neural progenitors¶

Markers from Fletcher et al. (2017):

- NEUROG1+, NEUROD1+

Markers from Nikaido:

- GAP43+

In [10]:

```
sc.pl.dotplot(
    adata,
    ["gap43", "neurog1", "neurod1", "stmn1b", "uchl1", "gng8"],
    "leiden",
    swap_axes=True,
)
```

In [11]:

```
markers[markers.gene.isin(["gap43", "neurog1", "neurod1", "stmn1b", "uchl1", "gng8"])]
```

Out[11]:

|  | cluster | gene | pval\_adj | LFC | pct.1 | pct.2 |
| --- | --- | --- | --- | --- | --- | --- |
| 220 | 0 | gng8 | 1.280569e-63 | 3.616665 | 10.330464 | 2.302366 |
| 7743 | 6 | gng8 | 6.942150e-20 | 2.351392 | 10.501355 | 2.302366 |
| 13903 | 10 | gap43 | 1.189273e-156 | 4.212159 | 58.914729 | 7.986018 |
| 14152 | 10 | uchl1 | 1.282342e-22 | 2.078373 | 16.623600 | 5.061845 |
| 14265 | 10 | stmn1b | 7.424579e-16 | 4.095423 | 7.751938 | 0.850363 |
| 14584 | 10 | neurod1 | 1.188067e-07 | 4.744189 | 3.359173 | 0.289056 |
| 14836 | 10 | neurog1 | 1.554705e-04 | 3.349569 | 2.153316 | 0.299140 |
| 15285 | 11 | gap43 | 1.473614e-45 | 2.453862 | 36.556604 | 7.986018 |
| 15408 | 11 | uchl1 | 7.083971e-20 | 2.093516 | 19.339623 | 5.061845 |
| 15643 | 11 | gng8 | 6.101139e-08 | 1.746865 | 9.080189 | 2.302366 |
| 15649 | 11 | stmn1b | 7.446858e-08 | 3.084925 | 5.660377 | 0.850363 |
| 16172 | 12 | gap43 | 7.359521e-50 | 2.791656 | 37.378641 | 7.986018 |
| 16369 | 12 | uchl1 | 2.857565e-11 | 1.780986 | 13.106796 | 5.061845 |
| 16715 | 12 | stmn1b | 1.144485e-03 | 2.068519 | 3.276699 | 0.850363 |
| 22364 | 16 | neurog1 | 6.419275e-07 | 5.586073 | 7.171315 | 0.299140 |
| 22617 | 16 | neurod1 | 2.957134e-05 | 4.264730 | 4.980080 | 0.289056 |
| 29207 | 20 | uchl1 | 4.307004e-07 | 1.500327 | 19.201995 | 5.061845 |
| 29581 | 21 | gng8 | 3.567676e-05 | 2.295858 | 11.356467 | 2.302366 |
| 33536 | 24 | uchl1 | 2.641331e-09 | 2.132084 | 27.888446 | 5.061845 |

Cluster 10 is INPs as it expresses both the progenitor markers neurog1+ and neurod1+ as well as the OSN marker gap43+. Also, it does not express gng8.

### Olfactory sensory neurons¶

OSNs are defined as UCHL1+, NST+, and OR+. They express CNG-family
genes in catfish. Tan et al. (2015) and Durante et al. (2020) make a distinction
between mature and immature OSNs, the latter based on GNG8 (immature) vs GNG13
(mature), so I've divided the markers into groups here based on whether they are
general or specific to mature or immature. GAP43 and OMP are two more classic
iOSN/mOSN markers, respectively (Huang et al., 2022).

Finally, Nikaido mentions a class of mOSNs that are also microvillar, defined by
gng8+, gap43-, trpc2b+, ompa-.

Some notes on orthology from Maggs:

- LOC103040890 is orthologous with mammalian gnal (not A mexicanus gnal)
- stmn1b is orthologous with mammalian stmn1 (not stmn1a)
- There is no A mexicanus ortholog with mammalian STOML3.

In [12]:

```
sc.pl.dotplot(
    adata,
    {
        "OSN": ["uchl1", "cnga2b", "cnga4"],
        "iOSN": ["gap43", "stmn1b"],
        "mOSN": ["LOC103040890", "gng13b", "ompa"],
        "mOSN-mv": ["gng8", "trpc2b"]
    },
    "leiden",
    swap_axes=True,
)
```

general OSN markers:

In [13]:

```
markers[markers.gene.isin(["uchl1", "cnga2b", "cnga4"])]
```

Out[13]:

|  | cluster | gene | pval\_adj | LFC | pct.1 | pct.2 |
| --- | --- | --- | --- | --- | --- | --- |
| 899 | 1 | cnga4 | 3.604553e-65 | 1.331309 | 25.376662 | 15.393923 |
| 927 | 1 | cnga2b | 3.247582e-45 | 1.577726 | 14.623338 | 7.740656 |
| 5034 | 3 | cnga4 | 6.868277e-65 | 1.434322 | 27.364621 | 15.393923 |
| 5056 | 3 | cnga2b | 9.222509e-51 | 1.799256 | 17.328520 | 7.740656 |
| 5832 | 4 | cnga4 | 1.072935e-57 | 1.257715 | 41.240046 | 15.393923 |
| 5886 | 4 | cnga2b | 3.243283e-39 | 1.505183 | 25.142207 | 7.740656 |
| 6775 | 5 | cnga4 | 3.098400e-39 | 1.371611 | 31.794195 | 15.393923 |
| 6924 | 5 | cnga2b | 3.829350e-15 | 1.174399 | 15.501319 | 7.740656 |
| 13476 | 9 | cnga2b | 4.274261e-20 | 1.389126 | 20.504202 | 7.740656 |
| 14152 | 10 | uchl1 | 1.282342e-22 | 2.078373 | 16.623600 | 5.061845 |
| 15408 | 11 | uchl1 | 7.083971e-20 | 2.093516 | 19.339623 | 5.061845 |
| 16369 | 12 | uchl1 | 2.857565e-11 | 1.780986 | 13.106796 | 5.061845 |
| 21376 | 15 | cnga4 | 1.009296e-09 | 1.056533 | 32.203390 | 15.393923 |
| 21456 | 15 | cnga2b | 4.480521e-06 | 1.233958 | 16.949153 | 7.740656 |
| 29207 | 20 | uchl1 | 4.307004e-07 | 1.500327 | 19.201995 | 5.061845 |
| 33287 | 24 | cnga4 | 1.688011e-27 | 2.065573 | 65.737052 | 15.393923 |
| 33536 | 24 | uchl1 | 2.641331e-09 | 2.132084 | 27.888446 | 5.061845 |
| 34401 | 25 | cnga4 | 4.424321e-08 | 1.520616 | 44.654088 | 15.393923 |

iOSN markers:

In [14]:

```
markers[markers.gene.isin(["gap43", "stmn1b"])]
```

Out[14]:

|  | cluster | gene | pval\_adj | LFC | pct.1 | pct.2 |
| --- | --- | --- | --- | --- | --- | --- |
| 13903 | 10 | gap43 | 1.189273e-156 | 4.212159 | 58.914729 | 7.986018 |
| 14265 | 10 | stmn1b | 7.424579e-16 | 4.095423 | 7.751938 | 0.850363 |
| 15285 | 11 | gap43 | 1.473614e-45 | 2.453862 | 36.556604 | 7.986018 |
| 15649 | 11 | stmn1b | 7.446858e-08 | 3.084925 | 5.660377 | 0.850363 |
| 16172 | 12 | gap43 | 7.359521e-50 | 2.791656 | 37.378641 | 7.986018 |
| 16715 | 12 | stmn1b | 1.144485e-03 | 2.068519 | 3.276699 | 0.850363 |

mOSN markers:

In [15]:

```
markers[markers.gene.isin(["LOC103040890", "gng13b", "ompa"])]
```

Out[15]:

|  | cluster | gene | pval\_adj | LFC | pct.1 | pct.2 |
| --- | --- | --- | --- | --- | --- | --- |
| 853 | 1 | ompa | 2.672064e-137 | 1.983353 | 32.850812 | 14.271309 |
| 5007 | 3 | ompa | 2.845201e-82 | 1.662164 | 29.494585 | 14.271309 |
| 5079 | 3 | gng13b | 5.853813e-43 | 1.559099 | 17.039711 | 8.577575 |
| 5743 | 4 | ompa | 3.564583e-175 | 2.294510 | 59.954494 | 14.271309 |
| 5796 | 4 | gng13b | 1.154734e-74 | 2.017448 | 33.333333 | 8.577575 |
| 6872 | 5 | gng13b | 2.437877e-19 | 1.316394 | 17.348285 | 8.577575 |
| 13365 | 9 | ompa | 7.258630e-58 | 1.737298 | 42.436975 | 14.271309 |
| 21378 | 15 | gng13b | 1.618257e-09 | 1.502006 | 21.280603 | 8.577575 |
| 33437 | 24 | gng13b | 2.808727e-12 | 1.875723 | 40.637450 | 8.577575 |
| 34471 | 25 | gng13b | 1.042875e-05 | 1.546071 | 30.188679 | 8.577575 |

mOSN-mv markers:

In [16]:

```
markers[markers.gene.isin(["gng8", "trpc2b"])]
```

Out[16]:

|  | cluster | gene | pval\_adj | LFC | pct.1 | pct.2 |
| --- | --- | --- | --- | --- | --- | --- |
| 7 | 0 | trpc2b | 0.000000e+00 | 4.173833 | 79.172452 | 18.126513 |
| 220 | 0 | gng8 | 1.280569e-63 | 3.616665 | 10.330464 | 2.302366 |
| 7420 | 6 | trpc2b | 0.000000e+00 | 3.135248 | 82.994580 | 18.126513 |
| 7743 | 6 | gng8 | 6.942150e-20 | 2.351392 | 10.501355 | 2.302366 |
| 15280 | 11 | trpc2b | 3.098716e-49 | 1.468604 | 55.542453 | 18.126513 |
| 15643 | 11 | gng8 | 6.101139e-08 | 1.746865 | 9.080189 | 2.302366 |
| 29023 | 20 | trpc2b | 6.004899e-48 | 1.755050 | 76.309227 | 18.126513 |
| 29390 | 21 | trpc2b | 4.408904e-53 | 2.643036 | 78.233438 | 18.126513 |
| 29581 | 21 | gng8 | 3.567676e-05 | 2.295858 | 11.356467 | 2.302366 |

Putting this all together:

- Clusters 0, 6, 20, and 21 are trpc2b+, gng8+, ompa-, indicating **microvillous mOSNs**.
- Clusters 1, 3, 4, 5, 9, 15, and 25 are gng13b+, ompa+, indicating **mOSNs**.
- Cluster 11 is gap43+ and stmn1b+ (immature), trpc2b+, gng8+, ompa- (microvillous), so this is **microvillous iOSNs**.
- Cluster 12 is gap43+ and stmn1b+ (immature), trpc2b-, gng8- (not microvillous), so this is **iOSNs**.
- Cluster 24 is cnga4+, uchl1+, gng13b+, ompa-, which is similar to OSNs but cannot properly be called OSNs without expressing ompa, so we will call these **OSN-like** for lack of a better description.

### Sustentacular cells¶

Sus cells are defined as SOX2+, PAX6+, HES1+. They are also JAG1+ and NOTCH2+,
but NOTCH1- and TP63-. This is a complicated pattern, so let's make a dot plot
with these markers grouped by whether they should be + or -. HES1 is the most
specific marker for Sus cells as SOX2 and PAX6 are also markers for HBC and GBC.

Notes on orthology:

- jag1a and jag1b are both orthologs of mammalian JAG1.
- only pax6a is an ortholog of mammalian PAX6
- only foxj1a is an ortholog of mammalian FOXJ1.
- her6 is the ortholog of mammalian HES1.

In [17]:

```
sc.pl.dotplot(
    adata,
    {
        "+": ["pax6a", "her6", "jag1a", "jag1b", "notch2", "sox2"],
        "-": ["notch1a", "notch1b", "tp63"],
        "ciliated": ["foxj1a"],
    },
    "leiden",
    swap_axes=True,
)
```

In [18]:

```
markers[
    markers.cluster.isin([2, 8, 13, 14, 22])
    & markers.gene.isin(
        ["pax6a", "her6", "jag1a", "jag1b", "notch2", "sox2", "dnah2", "foxj1a"]
    )
]
```

Out[18]:

|  | cluster | gene | pval\_adj | LFC | pct.1 | pct.2 |
| --- | --- | --- | --- | --- | --- | --- |
| 1732 | 2 | sox2 | 2.938943e-114 | 3.372988 | 24.056095 | 4.947567 |
| 2269 | 2 | jag1a | 2.443581e-39 | 2.807120 | 9.744696 | 2.430089 |
| 2426 | 2 | jag1b | 7.907329e-31 | 2.040004 | 9.996404 | 3.431702 |
| 2722 | 2 | pax6a | 3.759023e-21 | 1.711320 | 8.234448 | 3.290535 |
| 3677 | 2 | her6 | 1.004953e-07 | 2.149847 | 2.265372 | 0.742807 |
| 10745 | 8 | sox2 | 2.853380e-76 | 3.285533 | 32.565056 | 4.947567 |
| 10924 | 8 | jag1b | 5.423211e-49 | 3.413521 | 22.230483 | 3.431702 |
| 11372 | 8 | jag1a | 1.067659e-22 | 2.587068 | 12.193309 | 2.430089 |
| 11944 | 8 | pax6a | 2.518590e-10 | 1.501439 | 8.921933 | 3.290535 |
| 12072 | 8 | her6 | 5.786568e-09 | 2.963772 | 4.163569 | 0.742807 |
| 16884 | 13 | dnah2 | 1.443142e-298 | 3.558637 | 98.613518 | 29.487093 |
| 17481 | 13 | foxj1a | 8.087085e-26 | 6.647729 | 23.050260 | 0.628529 |
| 17508 | 13 | jag1b | 7.072199e-25 | 2.847682 | 28.422877 | 3.431702 |
| 20190 | 14 | sox2 | 2.878413e-16 | 2.005172 | 26.306620 | 4.947567 |
| 20699 | 14 | jag1a | 7.586734e-06 | 1.655549 | 10.627178 | 2.430089 |
| 20744 | 14 | jag1b | 1.606377e-05 | 1.414472 | 12.020906 | 3.431702 |
| 29897 | 22 | notch2 | 1.183026e-17 | 2.135093 | 42.307692 | 9.296854 |
| 30251 | 22 | jag1b | 1.928335e-07 | 2.686432 | 16.025641 | 3.431702 |

Based on this, it looks like 2, 8, 13, and 14 are sustentacular. 13 also has *dnah* family genes as 3 of its top 5 marker genes and expresses foxj1a, indicating that it has cilia. Triana-Garcia et al. (2021) differentiate between ciliated and secretory sustentacular cells in delta smelt, so I'm going to call 13 ciliated sustentacular and 2/8/14 secretory sustentacular.

### Microvillar cells¶

Microvillar cells are heterogenous in mammals, with distinct populations
characterized by TRPM5+, TRPC6+, or PLCB2+.

In [19]:

```
sc.pl.dotplot(
    adata, 
    ["trpm5", "plcb2", "trpc6a", "trpc6b"], 
    "leiden", 
    swap_axes=True
)
```

I'll call 27 TRPM5+ MCs and ?? TRPC6+ MCs. These clusters are too small for the
genes to be significant, but they are clearly expressed.

### Olfactory ensheathing glia¶

OEGs are NTFR+, S100+, VIM+, SOX10+ (Oprych et al., 2017).

Orthology:

- The antibodies Oprych used for S100 are probably generally reactive with many
  genes in the S100 family, so I'm plotting all of them that are expressed here.
- vim and viml are both orthologs of mammalian VIM
- ngfra and ngfrb are both orthologs of mammalian NGFR

In [20]:

```
sc.pl.dotplot(
    adata[adata.obs.leiden != "11"],
    ["s100u", "s100w", "s100a11", "s100a10b", "s100v2", "ngfra", "ngfrb", "apoc1", "vim", "viml", "sox10"],
    "leiden",
    swap_axes=True,
)
```

In [21]:

```
markers[markers.gene.isin(
    ["s100u", "s100w", "s100a11", "s100a10b", "s100v2", "ngfra", "ngfrb", "apoc1", "vim", "viml", "sox10"]
)]
```

Out[21]:

|  | cluster | gene | pval\_adj | LFC | pct.1 | pct.2 |
| --- | --- | --- | --- | --- | --- | --- |
| 2846 | 2 | s100u | 3.279434e-18 | 2.457702 | 5.142035 | 1.452003 |
| 3262 | 2 | s100a10b | 1.289205e-11 | 2.043464 | 3.595829 | 1.149503 |
| 3276 | 2 | s100w | 1.998875e-11 | 2.572469 | 2.912621 | 0.877252 |
| 3380 | 2 | s100v2 | 2.174544e-10 | 2.116918 | 3.056455 | 1.055391 |
| 4556 | 2 | sox10 | 1.418299e-03 | 3.781887 | 0.539374 | 0.087389 |
| 7159 | 5 | ngfra | 1.075785e-05 | 1.494788 | 4.155673 | 1.559559 |
| 7393 | 5 | vim | 7.825825e-03 | 1.886076 | 1.253298 | 0.406695 |
| 11828 | 8 | s100u | 1.018737e-11 | 2.447897 | 6.691450 | 1.452003 |
| 11886 | 8 | s100a10b | 4.533486e-11 | 2.701752 | 5.427509 | 1.149503 |
| 12467 | 8 | s100w | 7.988974e-06 | 2.160246 | 3.345725 | 0.877252 |
| 13113 | 8 | s100v2 | 3.396185e-03 | 1.286170 | 2.676580 | 1.055391 |
| 18434 | 13 | s100a10b | 1.550462e-08 | 2.720373 | 9.705373 | 1.149503 |
| 18747 | 13 | s100w | 2.293012e-06 | 2.446985 | 6.932409 | 0.877252 |
| 23706 | 16 | s100v2 | 9.734473e-03 | 2.162808 | 3.585657 | 1.055391 |
| 24151 | 17 | ngfrb | 7.155339e-14 | 4.531773 | 14.867617 | 0.910863 |
| 24375 | 17 | s100v2 | 9.997404e-10 | 3.818839 | 10.997963 | 1.055391 |
| 24601 | 17 | s100a10b | 2.771337e-07 | 3.405531 | 8.553971 | 1.149503 |
| 25465 | 17 | s100u | 4.009843e-03 | 1.683244 | 4.887984 | 1.452003 |
| 26439 | 18 | ngfrb | 1.677196e-07 | 3.973381 | 8.074534 | 0.910863 |
| 30101 | 22 | ngfrb | 1.642485e-09 | 4.412009 | 16.666667 | 0.910863 |
| 30298 | 22 | s100v2 | 6.998019e-07 | 3.615130 | 11.538462 | 1.055391 |
| 30794 | 22 | s100u | 2.845835e-04 | 2.776485 | 9.615385 | 1.452003 |
| 30913 | 22 | s100a10b | 7.077088e-04 | 2.808134 | 7.051282 | 1.149503 |
| 30936 | 22 | s100w | 8.481055e-04 | 3.069766 | 6.410256 | 0.877252 |
| 32131 | 23 | s100u | 3.567235e-09 | 3.672907 | 18.287938 | 1.452003 |
| 34915 | 26 | s100a11 | 5.870286e-09 | 7.880365 | 28.260870 | 0.373084 |
| 35408 | 26 | s100u | 8.586596e-04 | 3.287006 | 13.768116 | 1.452003 |
| 35456 | 26 | viml | 1.404496e-03 | 7.091921 | 10.144928 | 0.127723 |
| 36671 | 28 | s100u | 3.164014e-03 | 4.362575 | 17.910448 | 1.452003 |

I'm calling 26 OEGs because it's + for several s100 genes as well as viml. It's a tiny cluster, so it's not totally surprising that it's hard to figure out.

### Immune cells¶

In [22]:

```
sc.pl.dotplot(
    adata,
    {
        "leukocytes": ["ptprc"],
        "macrophages": ["mpp1", "c1qb", "cd4-1"],
        "T": ["lck"],
    },
    "leiden",
    swap_axes=True,
)
```

In [23]:

```
markers[markers.gene.isin(["ptprc", "mpp1", "c1qb", "cd4-1", "lck"])]
```

Out[23]:

|  | cluster | gene | pval\_adj | LFC | pct.1 | pct.2 |
| --- | --- | --- | --- | --- | --- | --- |
| 8183 | 7 | ptprc | 6.575753e-277 | 6.074219 | 65.745856 | 5.922291 |
| 8454 | 7 | cd4-1 | 2.013192e-38 | 4.056149 | 14.295580 | 1.929282 |
| 8662 | 7 | lck | 6.572001e-24 | 7.160356 | 7.734807 | 0.507529 |
| 9380 | 7 | mpp1 | 2.134427e-08 | 1.407568 | 6.698895 | 4.204759 |
| 21347 | 15 | ptprc | 5.712691e-13 | 1.540882 | 27.871940 | 5.922291 |
| 25710 | 18 | cd4-1 | 1.515698e-50 | 5.768562 | 45.548654 | 1.929282 |
| 25725 | 18 | mpp1 | 1.110563e-43 | 4.611071 | 42.236025 | 4.204759 |
| 25749 | 18 | ptprc | 1.130957e-37 | 3.038474 | 43.478261 | 5.922291 |
| 25809 | 18 | c1qb | 1.472627e-28 | 8.573028 | 26.293996 | 0.527696 |
| 27467 | 19 | ptprc | 1.380271e-41 | 3.643511 | 46.357616 | 5.922291 |
| 28043 | 19 | mpp1 | 7.433703e-07 | 1.985929 | 11.699779 | 4.204759 |
| 29502 | 21 | ptprc | 4.802280e-08 | 1.577384 | 25.867508 | 5.922291 |
| 29645 | 21 | cd4-1 | 3.573763e-04 | 1.837103 | 10.094637 | 1.929282 |
| 36668 | 28 | mpp1 | 3.077637e-03 | 2.883369 | 19.402985 | 4.204759 |

Based on this, I'm labelling 7 as T-cells, 18 as macrophages, and 19 as unspecified leukocytes.

### Otherwise unidentified cells¶

Cluster 22 has a weird mix of HBC (e.g., tp63+) and Sus (e.g., notch2+, jag1b+) markers. I will leave it as unknown unless anyone has any better ideas.

Cluster 23 is not yet identified.

In [24]:

```
markers[markers.cluster == 23].head(20)
```

Out[24]:

|  | cluster | gene | pval\_adj | LFC | pct.1 | pct.2 |
| --- | --- | --- | --- | --- | --- | --- |
| 31475 | 23 | dennd1b | 5.446368e-137 | 5.334140 | 98.054475 | 21.827104 |
| 31476 | 23 | mbnl1 | 1.326794e-129 | 4.487284 | 98.443580 | 20.922963 |
| 31477 | 23 | LOC125806044 | 2.071246e-121 | 8.408628 | 96.498054 | 3.240118 |
| 31478 | 23 | si:ch211-198m17.1 | 1.986139e-102 | 10.083043 | 94.552529 | 1.361253 |
| 31479 | 23 | fam49bb | 8.691205e-101 | 4.892022 | 94.941634 | 12.963834 |
| 31480 | 23 | LOC111195946 | 1.572646e-98 | 12.159231 | 91.439689 | 0.947835 |
| 31481 | 23 | LOC103028333 | 4.087959e-92 | 4.887372 | 92.996109 | 10.513579 |
| 31482 | 23 | igsf3 | 4.011799e-91 | 4.094738 | 94.163424 | 26.152864 |
| 31483 | 23 | dock1 | 1.746590e-90 | 4.158113 | 91.439689 | 17.686206 |
| 31484 | 23 | atp11a | 1.843458e-87 | 3.759085 | 92.217899 | 25.090750 |
| 31485 | 23 | acap3a | 1.289338e-83 | 6.642023 | 88.715953 | 3.771175 |
| 31486 | 23 | celf2 | 4.533720e-84 | 3.709615 | 92.217899 | 12.301694 |
| 31487 | 23 | pard3ba | 7.973188e-83 | 3.672957 | 91.828794 | 24.109304 |
| 31488 | 23 | diaph2 | 2.691078e-77 | 2.191522 | 97.276265 | 59.441382 |
| 31489 | 23 | nav3 | 9.999485e-77 | 3.448569 | 91.050584 | 25.662140 |
| 31490 | 23 | cd74a | 4.772616e-76 | 3.571160 | 92.217899 | 17.837456 |
| 31491 | 23 | sorl1 | 9.016199e-74 | 3.572886 | 89.105058 | 19.360043 |
| 31492 | 23 | si:dkey-220k22.1 | 1.922833e-73 | 5.376293 | 86.770428 | 5.754235 |
| 31493 | 23 | yrk | 6.722512e-69 | 7.140762 | 82.101167 | 1.851976 |
| 31494 | 23 | arhgap17b | 2.978772e-66 | 3.739408 | 83.657588 | 12.970557 |

In human data:

- dennd1b is a marker for astrocytes, and expressed in various neurons and glia
- mbnl1 has high expression in microglia and other glia, is a marker for
  oligodendrocytes
- fam49bb is an ortholog of CYRIB, which is expressed in various glia and is a
  marker for oligodendrocytes
- dock1 is a marker for oligodendrocytes
- atp11a is expressed in oligodendrocytes

So my best guess is oligodendrocytes.

Cluster 28 is not yet identified.

In [25]:

```
markers[markers.cluster == 28].head(20)
```

Out[25]:

|  | cluster | gene | pval\_adj | LFC | pct.1 | pct.2 |
| --- | --- | --- | --- | --- | --- | --- |
| 36427 | 28 | foxp1b | 1.589627e-39 | 5.521593 | 97.014925 | 18.788653 |
| 36428 | 28 | mctp1a | 2.567676e-34 | 7.545918 | 98.507463 | 4.056870 |
| 36429 | 28 | LOC111194333 | 1.364677e-27 | 14.351622 | 91.044776 | 0.252084 |
| 36430 | 28 | LOC103022645 | 1.950776e-24 | 11.085162 | 91.044776 | 0.789863 |
| 36431 | 28 | LOC103044987 | 7.190761e-23 | 10.247955 | 89.552239 | 0.883974 |
| 36432 | 28 | zfhx3b | 1.094996e-18 | 3.596089 | 83.582090 | 22.929551 |
| 36433 | 28 | LOC103022380 | 6.077860e-15 | 6.398133 | 71.641791 | 3.135924 |
| 36434 | 28 | fryl | 7.851539e-14 | 2.473798 | 80.597015 | 38.394058 |
| 36435 | 28 | LOC103042391 | 3.307434e-13 | 7.222270 | 62.686567 | 0.991530 |
| 36436 | 28 | tanc2a | 4.020667e-12 | 3.415772 | 67.164179 | 16.916510 |
| 36437 | 28 | LOC103027665 | 1.124628e-11 | 11.997877 | 61.194030 | 0.211750 |
| 36438 | 28 | itpr3 | 1.738800e-11 | 5.756186 | 59.701493 | 3.411535 |
| 36439 | 28 | LOC103025798 | 1.829306e-11 | 12.573399 | 56.716418 | 0.161334 |
| 36440 | 28 | bcas1 | 2.150307e-11 | 8.962944 | 59.701493 | 0.460473 |
| 36441 | 28 | wu:fj16a03 | 7.518651e-11 | 10.778322 | 55.223881 | 0.231917 |
| 36442 | 28 | flrt2 | 1.411320e-10 | 5.056125 | 53.731343 | 3.855203 |
| 36443 | 28 | sytl2b | 2.035526e-10 | 4.886191 | 58.208955 | 4.742538 |
| 36444 | 28 | LOC103024548 | 2.511452e-10 | 5.410859 | 53.731343 | 3.038451 |
| 36445 | 28 | kita | 7.127150e-10 | 6.397066 | 53.731343 | 2.006588 |
| 36446 | 28 | b4galt1 | 1.156208e-09 | 4.609240 | 53.731343 | 5.135789 |

No idea. Leaving as unknown.

### Final labels¶

Finally, let's apply the final labels to the adata object so that we can make a
nice plot. I've been keeping a separate spreadsheet, so I'm loading it here.

In [26]:

```
df = pd.read_csv("cell_type_assignments.csv")
df
```

Out[26]:

|  | Cluster | Cell type | Short name | Markers |
| --- | --- | --- | --- | --- |
| 0 | 0 | microvillous mature olfactory sensory neurons | mOSN-mv | gng8+, gap43-, trpc2b+, ompa- |
| 1 | 1 | mature olfactory sensory neurons | mOSN | gng13b+, ompa+, cnga4+, cnga2b+ |
| 2 | 2 | secretory sustentacular cells | sSus | sox2+, jag1a+, jag1b+, pax6a+, her6+, foxj1a-,... |
| 3 | 3 | mature olfactory sensory neurons | mOSN | gng13b+, ompa+, cnga4+, cnga2b+ |
| 4 | 4 | mature olfactory sensory neurons | mOSN | gng13b+, ompa+, cnga4+, cnga2b+ |
| 5 | 5 | mature olfactory sensory neurons | mOSN | gng13b+, ompa+, cnga4+, cnga2b+ |
| 6 | 6 | microvillous mature olfactory sensory neurons | mOSN-mv | gng8+, gap43-, trpc2b+, ompa- |
| 7 | 7 | T-cells | T-cells | ptprc+, cd4-1+, lck+, mpp1+ |
| 8 | 8 | secretory sustentacular cells | sSus | sox2+, jag1a+, jag1b+, pax6a+, her6+, foxj1a-,... |
| 9 | 9 | mature olfactory sensory neurons | mOSN | gng13b+, ompa+, cnga2b+ |
| 10 | 10 | immediate neural precursors | INP | gap43+, neurod1+, neurog1+, stmn1b+, uchl1+, g... |
| 11 | 11 | microvillous immature olfactory sensory neurons | iOSN-mv | gap43+, gng8+, trpc2b+, stmn1b+, uchl1+, ompa-... |
| 12 | 12 | immature olfactory sensory neurons | iOSN | gap43+, stmn1b+, uchl1+, gng8-, trpc2b- |
| 13 | 13 | ciliated sustentacular cells | cSus | jag1b+, notch2+, foxj1a+, dnah2+ |
| 14 | 14 | secretory sustentacular cells | sSus | sox2+, jag1a+, jag1b+, pax6a+, her6+, foxj1a-,... |
| 15 | 15 | mature olfactory sensory neurons | mOSN | gng13b+, ompa+, cnga4+, cnga2b+ |
| 16 | 16 | globose base cells | GBC | sox2+, pax6a+, six1b+, ascl1a+, neurod1+, neur... |
| 17 | 17 | horizontal base cells | HBC | tp63+, notch2+, notch1b+, sox2+, notch1a+, ceb... |
| 18 | 18 | macrophages | macrophages | ptprc+, cd4-1+, c1qb+, mpp1+ |
| 19 | 19 | leukocytes | leukocytes | ptprc+, mpp1+ |
| 20 | 20 | microvillous mature olfactory sensory neurons | mOSN-mv | gng8+, gap43-, trpc2b+, ompa- |
| 21 | 21 | microvillous mature olfactory sensory neurons | mOSN-mv | gng8+, gap43-, trpc2b+, ompa- |
| 22 | 22 | unknown | unknown (1) | NaN |
| 23 | 23 | oligodendrocytes | oligodendrocytes | dennd1b+, mbnl1+, fam49bb+, dock1+, atp11a+ |
| 24 | 24 | olfactory sensory neuron-like cells | OSN-like | cnga4+, uchl1+, gng13b+, ompa- |
| 25 | 25 | mature olfactory sensory neurons | mOSN | cnga4+, gng13b+, ompa+ |
| 26 | 26 | olfactory ensheathing glia | OEG | s100a11+, s100u+, viml+ |
| 27 | 27 | TRPM5+ microvillar cells | TRPM5+ MC | trpm5+ |
| 28 | 28 | unknown | unknown (2) | NaN |

In [27]:

```
df = pd.DataFrame(
    {
        "leiden": df["Cluster"].astype(str),
        "cell_type": df["Short name"].astype("category")
    }
)
adata.obs = pd.merge(adata.obs.reset_index(), df).set_index("index")
```

In [28]:

```
sc.pl.umap(adata, color="cell_type", legend_loc="on data", save="_cluster_names.pdf")
```

```
WARNING: saving figure to file figures/umap_cluster_names.pdf
```

In [29]:

```
adata.write_h5ad("clusters_labeled.h5ad")
```

### Works cited¶

Bryche, B., Baly, C., & Meunier, N. (2021). Modulation of olfactory signal detection in the olfactory epithelium: focus on the internal and external environment, and the emerging role of the immune system. *Cell and Tissue Research*, 384(3), 589–605.

Calvo-Ochoa, E., Byrd-Jacobs, C. A., & Fuss, S. H. (2021). Diving into the streams and waves of constitutive and regenerative olfactory neurogenesis: insights from zebrafish. *Cell and Tissue Research*, 383(1), 227–253.

Chen, M., Reed, R. R., & Lane, A. P. (2019). Chronic Inflammation Directs an Olfactory Stem Cell Functional Switch from Neuroregeneration to Immune Defense. *Cell Stem Cell*, 25(4), 501–513.e5.

Durante, M. A., Kurtenbach, S., Sargi, Z. B., Harbour, J. W., Choi, R., Kurtenbach, S., Goss, G. M., Matsunami, H., & Goldstein, B. J. (2020). Single-cell analysis of olfactory neurogenesis and differentiation in adult humans. *Nature Neuroscience*, 23(3), 323–326.

Fletcher, R. B., Das, D., Gadye, L., Street, K. N., Baudhuin, A., Wagner, A., Cole, M. B., Flores, Q., Choi, Y. G., Yosef, N., Purdom, E., Dudoit, S., Risso, D., & Ngai, J. (2017). Deconstructing Olfactory Stem Cell Trajectories at Single-Cell Resolution. *Cell Stem Cell*, 20(6), 817–830.e8.

Fritzch, B., & Elliot, K. L. (2022). The Senses: Perspectives from Brain, Sensory Ganglia, and Sensory Cell Development in Vertebrates. In B. Fritzch & K. L. Elliot (Eds.), *Evolution of Neurosensory Cells and Systems* (pp. 1–27). Taylor & Francis.

Genovese, F., & Tizzano, M. (2018). Microvillous cells in the olfactory epithelium express elements of the solitary chemosensory cell transduction signaling cascade. *PloS One*, 13(9), e0202754.

Getchell, M. L., & Getchell, T. V. (1992). Fine structural aspects of secretion and extrinsic innervation in the olfactory mucosa. Microscopy Research and Technique, 23(2), 111–127.
Goss, G. M., Chaudhari, N., Hare, J. M., Nwojo, R., Seidler, B., Saur, D., & Goldstein, B. J. (2016). Differentiation potential of individual olfactory c-Kit+ progenitors determined via multicolor lineage tracing. *Developmental Neurobiology*, 76(3), 241–251.

Goulding, E. H., Ngai, J., Kramer, R. H., Colicos, S., Axel, R., Siegelbaum, S. A., & Chess, A. (1992). Molecular cloning and single-channel properties of the cyclic nucleotide-gated channel from catfish olfactory neurons. *Neuron*, 8(1), 45–58.

Hanchate, N. K., Kondoh, K., Lu, Z., Kuang, D., Ye, X., Qiu, X., Pachter, L., Trapnell, C., & Buck, L. B. (2015). Single-cell transcriptomics reveals receptor transformations during olfactory neurogenesis. *Science*, 350(6265), 1251–1255.

Huang, J. S., Kunkhyen, T., Rangel, A. N., Brechbill, T. R., Gregory, J. D., Winson-Bushby, E. D., Liu, B., Avon, J. T., Muggleton, R. J., & Cheetham, C. E. J. (2022). Immature olfactory sensory neurons provide behaviourally relevant sensory input to the olfactory bulb. *Nature Communications*, 13(1), 6194.

Kocagöz, Y., Demirler, M. C., Eski, S. E., Güler, K., Dokuzluoglu, Z., & Fuss, S. H. (2022). Disparate progenitor cell populations contribute to maintenance and repair neurogenesis in the zebrafish olfactory epithelium. *Cell and Tissue Research*, 388(2), 331–358.

Lazzari, M., Bettini, S., & Franceschini, V. (2013). Immunocytochemical characterization of olfactory ensheathing cells in fish. *Brain Structure & Function*, 218(2), 539–549.

Oprych, K., Cotfas, D., & Choi, D. (2017). Common olfactory ensheathing glial markers in the developing human olfactory system. *Brain Structure & Function*, 222(4), 1877–1895.

Schwob, J. E., Jang, W., Holbrook, E. H., Lin, B., Herrick, D. B., Peterson, J. N., & Hewitt Coleman, J. (2017). Stem and progenitor cells of the mammalian olfactory epithelium: Taking poietic license. *The Journal of Comparative Neurology*, 525(4), 1034–1054.

Tan, L., Li, Q., & Xie, X. S. (2015). Olfactory sensory neurons transiently express multiple olfactory receptors during development. Molecular Systems Biology, 11(12), 844.
Triana-Garcia, P. A., Nevitt, G. A., Pesavento, J. B., & Teh, S. J. (2021). Gross morphology, histology, and ultrastructure of the olfactory rosette of a critically endangered indicator species, the Delta Smelt, Hypomesus transpacificus. *Journal of Comparative Physiology. A, Neuroethology, Sensory, Neural, and Behavioral Physiology*, 207(5), 597–616.
